# Evidence for managing herbivores for reef resilience

**DOI:** 10.1101/2023.09.19.557828

**Authors:** Mary K. Donovan, Chelsie W.W. Counsell, Megan J. Donahue, Joey Lecky, Laura Gajdzik, Stacia D. Marcoux, Russell Sparks, Christopher Teague

**Affiliations:** Hawai‘i Monitoring and Reporting Collaborative (HIMARC); Center for Global Discovery and Conservation Science, School of Geographical Sciences and Urban Planning, Arizona State University; Cooperative Institute for Marine and Atmospheric Research; Hawai‘i Institute of Marine Biology, University of Hawai‘i at Mānoa; Pacific Islands Regional Office, National Oceanic and Atmospheric Administration; Division of Aquatic Resources, Department of Land and Natural Resources, State of Hawai‘i, USA

**Keywords:** resilience-based management, coral, human impacts, Hawai‘i

## Abstract

Herbivore management is an important tool for resilience-based approaches to coral reef conservation. Yet, evidence-based science is needed to enact successful management. We synthesized data from multiple monitoring programs in Hawai’i to measure herbivore biomass and benthic condition over a 10-year period preceding any major coral bleaching. We analyzed data from 20,242 transects alongside data on 27 biophysical and human drivers and found herbivore biomass was highly variable throughout Hawai’i, with high values in remote locations and the lowest values near population centers. Both human and biophysical drivers explained variation in herbivore biomass, and among the human drivers both fishing and land-based pollution had negative effects on biomass. We also found evidence that herbivore functional group biomass is strongly linked to benthic condition, and that benthic condition is sensitive to changes in herbivore biomass associated with fishing. We show that when herbivore biomass is below 80% of potential biomass benthic condition is predicted to decline. We also show that a range of management actions, including area-specific fisheries regulations and gear restrictions, can increase parrotfish biomass. Together, these results provide lines of evidence to support managing herbivores as an effective strategy for maintaining or bolstering reef resilience in a changing climate.

## Introduction

Coral reefs worldwide are threatened by increasingly severe local and global stressors [1,2]. The ultimate solution to limiting negative social and ecological effects that stem from climate-driven coral loss is concerted global action on climate change. However, local action is also needed in the near-term to mitigate reef decline from direct adverse effects of local stressors [3,4] and interactions with heat stress that exacerbate the effects of climate change [5,6]. Resilience-based management is one opportunity for local-management action aimed at maintaining or increasing the resistance and/or recovery capacity of reefs through reduction of local stressors and/or the maintenance or bolstering of key ecological processes to promote ecosystem resistance and recovery from heat stress [7–9].

While there are multiple avenues that may be taken under the umbrella of resilience-based management [8–10], managing for herbivory by herbivorous fishes has been proposed as an especially effective option [11–13]. Herbivory contributes to maintaining a balance between corals and algae on reefs and facilitates coral recovery and recruitment [14,15]. The goal of herbivore-based management is thus to maintain sufficient levels of herbivory through actions to either restore or conserve populations of herbivorous species [16,17], while also considering the cultural and food security aspects of herbivore fisheries [18].

Managing herbivores with the overarching goal of increasing coral reef resilience relies on several assumptions [17]. First is the assumption that increased herbivory alters benthic condition in a way that is favorable for corals. Experiments have consistently shown that reducing herbivory leads to increases in algal biomass and canopy height [15,19,20]. Similarly, spatial associations between herbivore biomass and macroalgal cover have been observed in multiple regions, across both Caribbean and Pacific basins [13,21–23], although herbivore-benthic patterns are known to vary across these regions [24]. The relationship between fishes and the benthos is complex, reciprocal, and operates on multiple scales, so successful herbivore-based management will be context-dependent and likely related to the relative distribution of different functional groups within the herbivore guild [25]. A second assumption of herbivore-based management is that management actions (e.g., restrictions on fishing) can influence herbivory by increasing the abundance, size, and diversity of herbivores [17]. Demonstrating a management effect depends on measurements of the baseline conditions of coral and herbivore populations prior to implementation of management actions.

Herbivore-based management was recently identified as a priority for resilience-based management by the State of Hawai’i following severe coral bleaching in 2014-15 that resulted in 50% loss of corals in some areas [26], and a systematic review of resilience-based strategies that emphasized herbivore-based management practices [10]. Effective implementation of herbivore-based management as a tool for improving reef resilience requires an evaluation of underlying assumptions in a local ecosystem context: how increasing herbivory will influence benthic condition and which management actions can influence herbivore populations.

Combining a decade of fish and benthic surveys from multiple institutions and 27 biophysical and human data layers in Hawai’i we assessed reef resilience as benthic condition measured as the log-ratio of calcified to macroalgal cover and its relationship to herbivores and other drivers. Our study aimed to answer the following questions to inform herbivore-based management: (1) what biophysical and human factors influence herbivorous fishes, (2) what are spatial patterns of herbivorous fishes at scales relevant to management, (3) is there evidence that benthic condition is related to herbivorous fishes, (4) does fishing as a driver of herbivorous fishes influence benthic condition, and (5) is there evidence that herbivore-based management improves herbivore populations? Together, addressing these questions provides evidence to support implementing resilience-based management by bolstering herbivory in reef systems.

## Results

### Drivers of herbivore assemblages

A range of drivers influenced total herbivore biomass (Fig 1), including metrics of fishing, land-based pollution, oceanography, and habitat. Among the human drivers, non-commercial boat-based spearfishing and total boat-based net fishing had the largest negative effects on total herbivore biomass, followed by urban runoff and on-site waste disposal effluent. Biophysical drivers also explained variability in total herbivore biomass: greater and more frequent wave anomalies, higher irradiance, less frequent chlorophyll-*a* anomalies, and lower temperature variation were associated with higher total herbivore biomass. Habitat characteristics were among the strongest drivers of herbivore biomass with greater herbivore biomass associated with higher benthic rugosity, shallower depths, and reef habitat.

**Figure 1.**
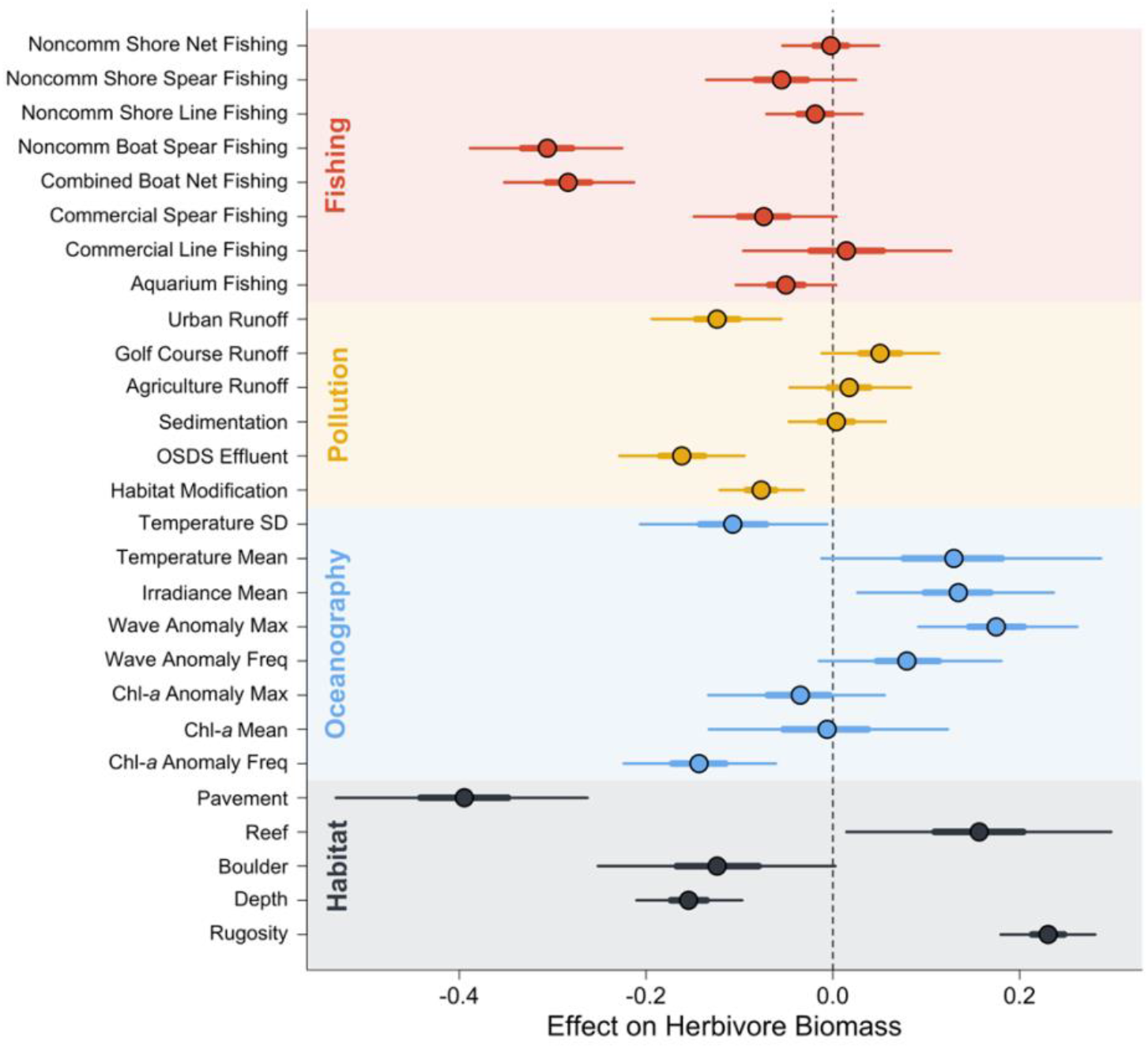
Coefficients from a Bayesian Hierarchical model estimating the relationship between total herbivore biomass and drivers that include metrics of fishing (red), land-based pollution (yellow), oceanography (blue), and habitat (grey). Circles are medians and horizontal lines correspond to 50% intervals (thick lines) and 95% intervals (thin lines) of posterior distributions of each coefficient. We interpret evidence of a relationship between the driver and herbivore biomass if either or both intervals do not overlap zero. Noncomm = non-commercial fishing; OSDS = on-site waste disposal systems; SD = standard deviation; Max = maximum; Freq = frequency; Chl-*a* = chlorophyll-*a*.

Considering the three herbivore functional groups separately (i.e., browsers, grazers, and scrapers), patterns between biomass and driver variables were generally consistent (Fig S1). Browser biomass had a greater negative association with on-site waste disposal effluent and habitat modification, and a less negative association with non-commercial boat-based spearfishing than other functional groups. Browser biomass was more strongly positively associated with maximum wave anomalies and more strongly negatively associated with depth than other functional groups. Grazer biomass was less strongly related to rugosity and scraper biomass was more strongly related to rugosity than overall herbivore biomass.

### Spatial patterns of herbivore assemblages

Herbivore biomass varied spatially across Hawai’i (Fig S2), with post-stratified mean biomass by moku (traditional land divisions) ranging between 7.3 and 158.7 g m^-2^ (Fig 2a). The highest biomass values were observed in remote areas, including moku on the islands Kaho’olawe and Ni’ihau, the north shore of Moloka’i, the Nāpali coast of Kaua’i, the Kahikinui and Hāna coastlines of Maui, and the Hāmākua and Ka’ū coastlines of Hawai’i island (Fig 2a, Fig S2). Herbivore biomass was generally lower around O’ahu (Fig 2a, Fig S2), with post-stratified mean values of all O’ahu moku falling in the lowest quartile of moku statewide (Fig 2a). Biomass in 19 of 43 moku were lower than biomass inside no-take marine reserves when comparing 50% posterior intervals (Fig S3).

**Figure 2.**
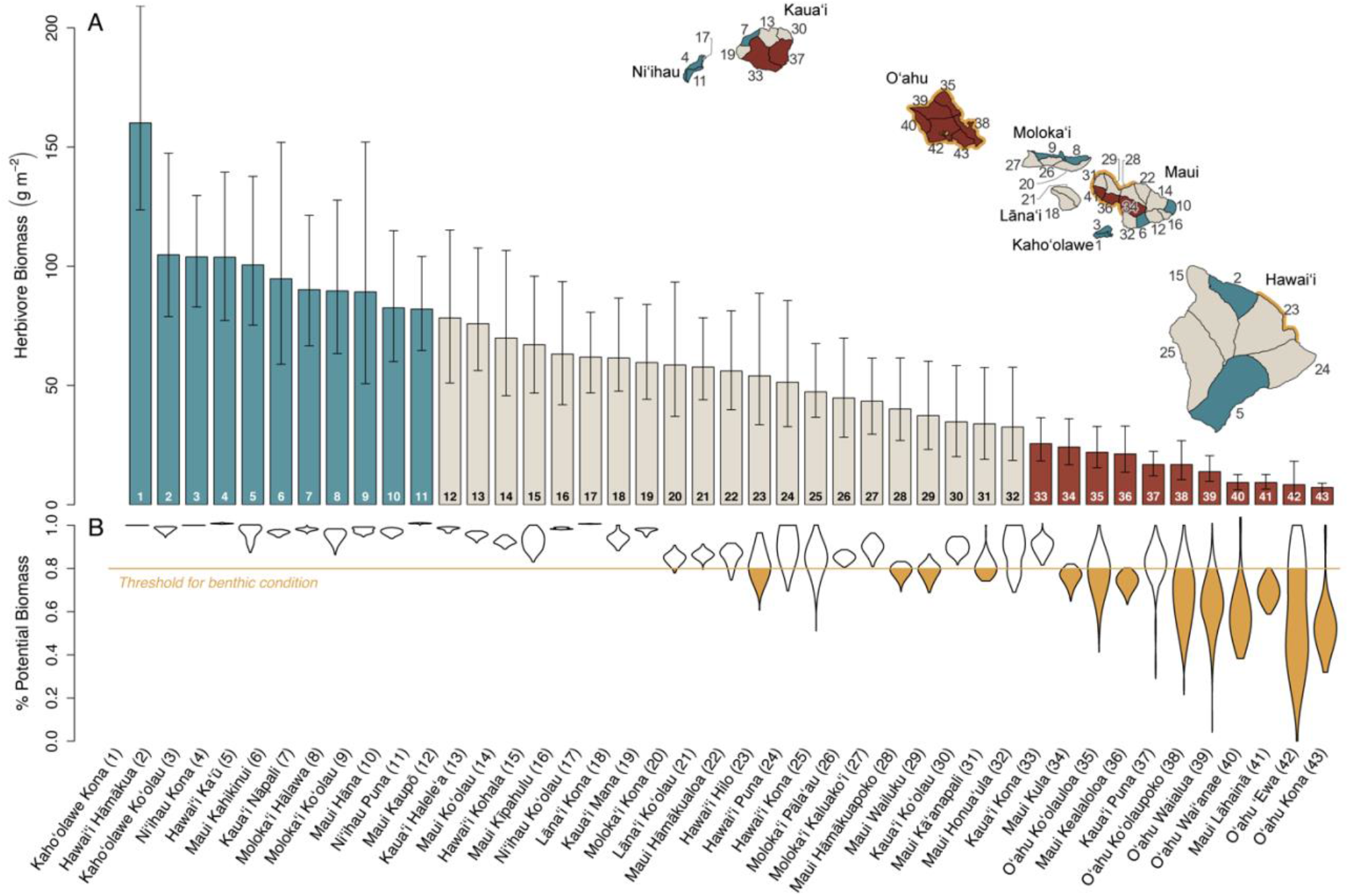
(A) Posterior estimates of herbivore biomass summarized across moku (land divisions). Bars are means and error bars are 50% intervals of posterior predictions for all 100 m^2^ pixels in each moku, and thus are post-stratified estimates that account for the relative distribution of habitat and variation in other predictors. Moku with mean values in the upper quartile (top fourth) are colored turquoise, and moku with mean values in the lower quartile (bottom forth) are colored red. Numbers correspond to labels at the bottom of (B) and to numbers on map (inset) (a larger map of moku boundaries with labels is provided in Fig S5). (B) Violin plots of percent potential biomass for all 100 m^2^ pixels in each moku estimated as the expected biomass from model in Fig 1 divided by the potential biomass when fishing is minimized and all other predictors are held constant, with threshold (80%) below which benthic condition is predicted to decrease (from Fig 3b). Moku with >44% of area under the threshold are colored yellow in B and the coastline is colored yellow in the inset.

### Benthic-driver relationships

Multiple drivers influenced spatial variability in benthic condition, measured as the log-ratio of calcified (coral + crustose coralline algae) and macroalgal cover, together explaining 51% of overall variability. Biomass of scrapers, grazers, and browsers and two- and three-way interactions between them were strongly associated with benthic condition (Fig 3a), compared to other drivers. Visualization of marginal effects across the three groups revealed several patterns (Fig S4). The highest levels of the benthic condition ratio (greater calcified cover relative to macroalgal cover) were associated with high biomass of either abundant scrapers or with high combined grazers and browsers. Higher macroalgal cover relative to calcified cover was associated with low biomass of all three functional groups.

**Figure 3.**
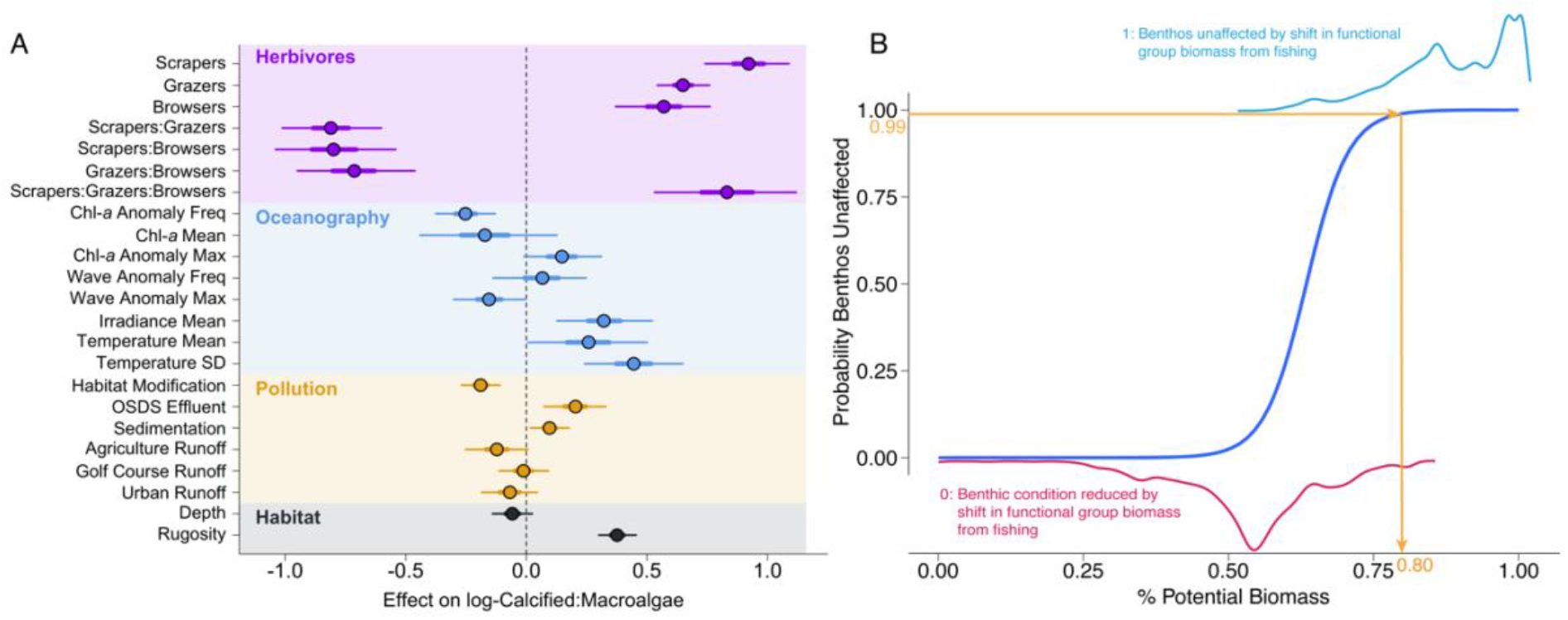
Relationship between herbivore functional group biomass and benthic condition. (A) coefficients from a Bayesian Hierarchical model estimating the relationship between biomass of three herbivore functional groups (scrapers, grazers, browsers), measures of oceanographic conditions, pollution, and depth and rugosity as predictors and the log-ratio of calcified benthic cover (coral + coralline algae) and macroalgal cover. Circles are medians and horizontal lines correspond to 50% intervals (thick lines) and 95% intervals (thin lines) of posterior distributions of each coefficient. (B) Probability from logistic regression (dark blue line) that benthic condition is unaffected by difference in herbivore biomass between expected and potential levels with fishing minimized as a function of percent of potential herbivore biomass (expected/potential) for each 100m^2^ pixel in the study domain. Pixels were classified as either unaffected (1) or affected (log(calcified:macroalgae) increased) (0), with density distributions of all % potential biomass values plotted above (light blue) and below (magenta), respectively. Yellow horizontal and vertical lines show where the probability of benthic change of 0.99 (y-axis) corresponds to % potential herbivore biomass (x-axis = 80%).

In addition to herbivores, other drivers helped explain variation in benthic condition, including oceanographic, pollution, and rugosity drivers (Fig 3a). Lower anomalies and higher maximum chlorophyll-*a* values, lower wave anomaly maximums, and higher mean irradiance, mean temperature, and standard deviations of temperature, as well as higher rugosity were all associated with greater calcified cover relative to macroalgal cover. The effects of different measures of land-based pollution were mixed with negative effects of habitat modification and agricultural and urban runoff, and positive effects of waste disposal effluent and sediment. Given the evidence for positive effects of effluent and sediment were counter to the hypothesized direction of these effects, we explored each pattern further. 70% of the top 10% of observations of waste-disposal effluent were from a single no-take area, Pūpūkea on the north shore of O’ahu. To evaluate the leverage of these observations, we set the effluent values for those observations to zero and re-ran the model and found that the effect of effluent was no longer distinguishable from zero. Similarly, 31% of the replicates with high values of sediment and high benthic condition were from Kohala on Hawai’i island, and there was no evidence of a sediment effect without those observations.

### Patterns of fishing effects on herbivores and benthos

After accounting for spatial variation in other drivers, we found that there was a 99% probability that benthic condition is not below expected condition when the biomass of herbivores is greater than 80% of potential biomass (Fig 3b). Across our study domain (Fig S2), 30% of all pixels are below this threshold, 44%-99% of the within-moku area was below this threshold in 13 of 43 moku, including all O’ahu moku, with 4 other moku <11 (Fig 2b). The entire area of 23 of 40 moku, largely remote areas with relatively high herbivore biomass, were above the 80% threshold (Fig 2a,b). In general herbivore biomass and the distribution of % potential biomass scaled with one another, with a few exceptions. For example, the entire area of the Kaua’i Kona moku was above the threshold despite being in the lowest quantile of biomass, indicating other drivers besides fishing are limiting biomass in that moku.

### Management effectiveness for herbivores

Multiple spatial- and gear-based actions have been effective in increasing the biomass of herbivores in Hawai’i based on analyses that combined measures of presence/absence and biomass when present using binomial-Gamma hurdle models (Fig 4, Fig S5). Across the island of Maui, the probability of observing parrotfish was significantly higher inside three no-take marine reserves compared to eight non-reserve sites monitored in 2018 and 2019 (df = 1,987, z = 7.64, p < 0.01), but parrotfish biomass when present was not significantly different between marine reserve and non-reserve sites (df = 1,583, t = 0.16, p = 0.88) (Fig 4a, Table S1). At Kahekili, a reef located on the northwestern coast of Maui, both the probability of observing parrotfish and parrotfish biomass when present were higher in five years following a prohibition on the take of herbivorous fishes, compared to the two years before the closure (Fig 4b, Table S1), with the biomass 1.1 times higher after the closure (df = 1,931, t = 2.06, p = 0.040). On the west coast of Hawai’i island a ban on SCUBA spearfishing was implemented in 2013, and a comparison of 10 sites outside of no-take reserves between 2007-2010 and 2016-2019 resulted in increasing probability of observing parrotfish (df = 1,309, z = 1.99, p = 0.046), and there was a marginally significant (α < 0.10) increase in the biomass when present (df = 1,282, t = 1.71, p = 0.089) (Fig 4c, Table S1). Across 13 fisheries management areas (which restrict aquarium take among other gear restrictions) there was no difference in the probability of observing parrotfish, and a marginally significant increase in biomass when present (df = 1,376, t = 1.78, p = 0.076). Inside no-take reserves parrotfish biomass increased significantly between the same time periods with 1.2 times higher biomass after the spearfishing ban than before (df = 1,58, t = 2.75, p = 0.008).

**Figure 4.**
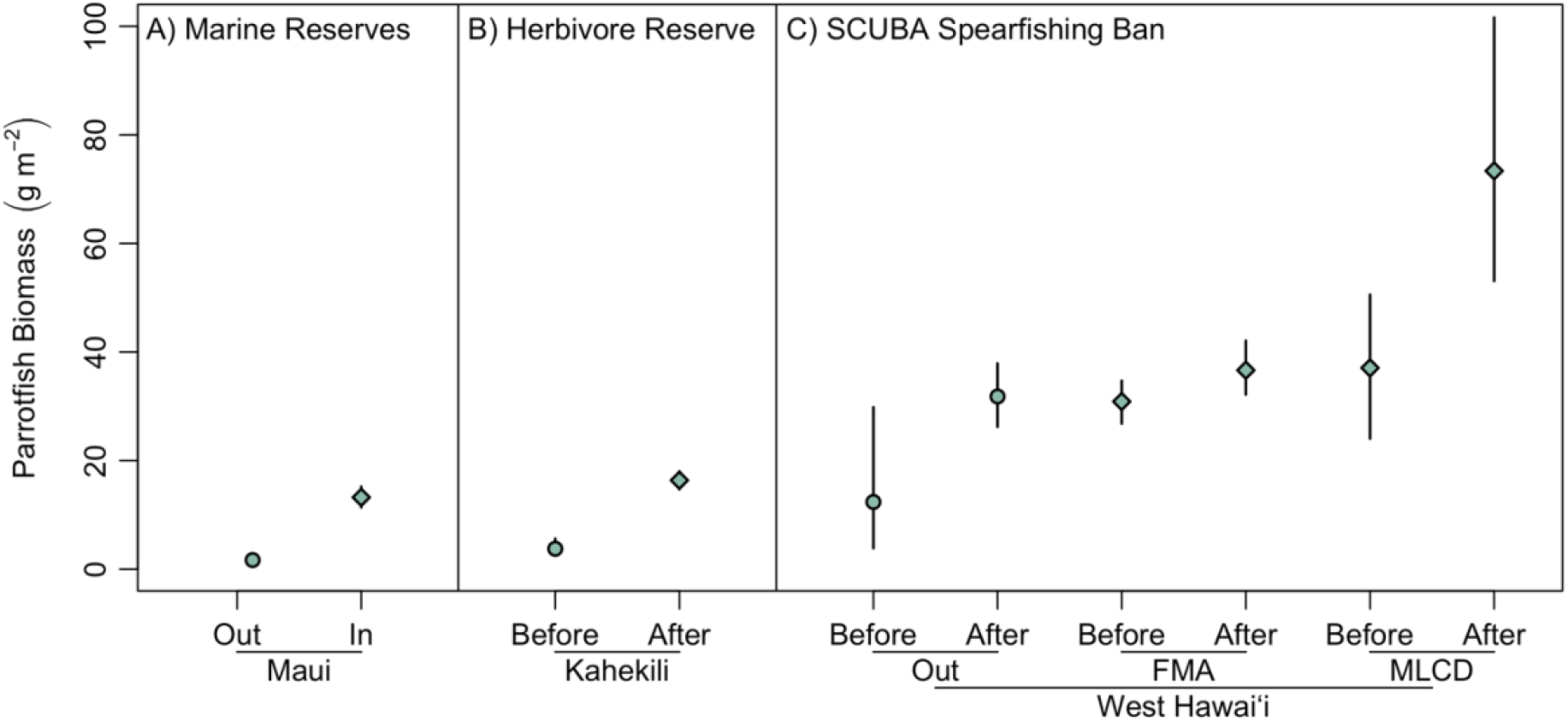
Trends in parrotfish biomass across management interventions at three locations in Hawai’i. Shapes are means and lines are 95% intervals from bootstrapped predictions from a binomial-Gamma hurdle model that combine predictions of probability of presence (binomial component) and biomass when present (Gamma component). (A) Comparison of 3 marine reserves (circle), and 8 non-reserve sites (diamond) on the island of Maui based on surveys conducted in 2018 and 2019. (B) Comparison before and after establishment of Kahekili Herbivore Fisheries Management Area on Maui established in 2009, before data are from 2008 and 2009, and after data from 2011-2015. (C) Comparison of sites in the West Hawai’i Fisheries Management Area before and after a ban on SCUBA spearfishing took effect in 2013 with before data from 2007-2010 and after data from 2016-2019, with comparisons split between 10 sites not in protected areas (“Out”), 13 sites inside Fishery Management Areas (aquarium collecting prohibited and other gear restrictions in some areas, “FMA”), and 2 sites in Marine Life Conservation Districts (no-take areas, “MLCD”). There were few or no observations without parrotfishes inside the no-take areas in West Hawai’i so means and intervals are from a Gamma model rather than a hurdle model for the “MLCD” portion of (C).

## Discussion

Using high resolution, broad scale spatial datasets on reef condition and human and biophysical drivers, we provide a basis for herbivore-based management in Hawai’i and a case study for management in other regions. We show that herbivore biomass is influenced by fishing, land-based pollution, and biophysical variables. By measuring or down-scaling this wide range of drivers at a fine spatial scale (100 m^2^), our results provide estimates of herbivore populations at scales that are relevant to management. We found strong associations between the biomass of herbivore functional groups and benthic condition, supporting existing evidence that some spatial variability in reef condition can be attributed to herbivores. We modeled how herbivore biomass was influenced by fishing in the context of other drivers and combined those predictions with models of benthic condition. We found that pixels fished below 80% of their pixel-specific potential herbivore biomass were sufficiently reduced from fishing to be associated with decreased benthic condition. Further, we found evidence for the effectiveness of multiple herbivore management approaches, confirming that these fisheries interventions can maintain and increase herbivore presence and biomass. Together, these results provide evidence that managing herbivore biomass can play a role in mediating benthic condition, which could support maintaining or bolstering reef resilience.

Our high-resolution predictions of herbivore biomass reveal important spatial features, including particularly high herbivore biomass in remote areas and low herbivore biomass in areas that are accessible and with large human populations (Fig 2, Fig S2). These patterns are similar to those that have been observed previously for total fish biomass and resource fish biomass in Hawai’i [27,28], and elsewhere [29,30]. We have extended this understanding by incorporating fine-scale variation of both human and environmental variables. Large bio-physical gradients exist in Hawai’i across important structuring variables such as wave energy, depth, and rugosity [31]. Thus, our ability to represent spatial variability in ecosystem metrics for management, such as herbivore biomass, is dependent on our ability to incorporate variation in these highly influential gradients [32]. To produce reliable estimates of biomass associated with particular geographies that are relevant for management (i.e., islands, moku, reserves), we post-stratified our estimates of ecosystem metrics to represent the spatial variability in the underlying biophysical conditions within each geography of interest. Thus, our results represent estimates that incorporate important underlying variability in human and environmental drivers, which overcome limitations associated with unbalanced sampling when combining data from multiple sources and provides the best available biomass predictions for informing management decisions.

We show clear correlational links between herbivore functional group biomass and benthic condition based on an extensive dataset that included variation due to other drivers. The combined effects of the three herbivore functional groups were complex, with strong two- and three-way interactions in relationship to log-ratio of calcified and macroalgal cover (Fig 3a, Fig S4). These interactions reflect the distinct and complementary functional roles each group plays in maintaining the balance between calcifiers and macroalgae [25]. All three main effects were positively associated with a calcified benthic state, reflecting the role each herbivore functional group plays in consuming algae. The groups also complement each other by performing that role differently, which was reflected in the negative two-way interactions and positive three-way interaction. For example, browsers and scrapers are often associated with contrasting benthic states [33]. Our results followed this pattern, with calcified cover maximized relative to macroalgae when browsers are absent, and scrapers are in high abundance (Fig S4). Grazers also played a strong role in differentiating between benthic states especially in context of the other two functional groups, which may relate to the ubiquity of grazers in Hawai’i compared to other regions [34,35]. We also found that habitat was a strong driver of herbivore biomass (Fig 1, Fig S1), further emphasizing the complex reciprocal relationship between the two components of the system [36,37]. Untangling these complex relationships among functional groups and benthic condition further is an important next step for setting resilience-based management targets.

Multiple types of fishing were negatively associated with herbivore biomass, with spearfishing and boat-based net fishing (both commercial and non-commercial) having particularly strong effects. We also reviewed evidence from fisheries interventions, which demonstrate that multiple approaches are available to bolster parrotfish populations. Herbivorous fishes are important components of subsistence and culture in island communities like Hawai’i. Given this, opportunities for management should be sought that consider equitable pathways recognizing the diverse needs of local communities given the potential for successful outcomes with partial protections that can complement or replace total closures [38]. Our results support potential compromises for fisheries management that can focus on specific functional groups of reef fishes (e.g., parrotfishes), specific locations (e.g., Kahekili), or specific gears (e.g., limits to SCUBA spearfishing). However, the relative effectiveness of area-based and gear-based actions was not clear in our analyses given that biomass increased both inside and outside of marine reserves in West Hawai’i after a ban on SCUBA spearfishing (Fig 4c). The success of herbivore-based management can be bolstered by careful consideration of the appropriate sizing and spacing of marine reserves [39], adequate information on species-specific life history and population assessments for species-specific actions, support and capacity for enforcement within and across locations, and needs and rights of indigenous and local communities [38,40].

We uncovered large variability in herbivore biomass across Hawai’i (Fig 2, Fig S2), with total herbivore biomass in 44% of moku below the mean biomass in no-take marine reserves (Fig S3), and parrotfish biomass within reserves was much lower across Maui than West Hawai’i (Fig 4), bringing to question what the appropriate herbivore biomass reference is for maintaining reef resilience, defined here as higher calcified cover relative to macroalgae. The case studies we investigated here are limited to changes in herbivore biomass, so we are unable to assess how fisheries management translates to changes in the benthos in our case studies. However, a previous study from Kahekili found evidence for increases in calcified cover following increases in herbivores post-closure [41], indicating the possibility for herbivore management to increase benthic resilience for Hawai’i reefs. We also investigated the role of fishing of herbivores on the benthos by combining results from our model of drivers of herbivore biomass with our model of benthic condition to uncover a threshold of 80% of potential herbivore biomass, below which benthic condition is predicted to decline (Fig 3b). We calculated percent potential biomass as expected biomass given measured fishing levels divided by the potential biomass without fishing while accounting for spatial variation in other drivers. Thus, we were able to uncover the sensitivity of the benthos to fishing effects on biomass of herbivore functional groups, and how pervasive those negative effects were across the study domain. This threshold aligns with previous work from other regions that have suggested a ‘safe operating space’ for fish biomass, below which algae populations are predicted to proliferate [42–44], including a theoretical model based on Caribbean fisheries that suggested a similar threshold of 90% [45]. We found that % potential biomass for 30% of the study domain was below the 80% threshold, and that much of that area was in more populated and more accessible areas (Fig 2). These results underscore not only the potential for compromised cultural and food security that these fisheries provide, but also that those declines in herbivore biomass are associated with a compromised benthos that could lead to further negative impacts for the ecosystem services that reefs provide to the people of Hawai’i.

Our results also show negative effects of land-based pollution, in addition to fishing effects, on herbivore biomass, emphasizing that fisheries management is not a panacea. We find that herbivore biomass is lower in areas with increased urban runoff, cesspool effluent, and coastal modifications (e.g., seawalls). The mechanisms linking these land-based drivers to herbivore biomass are not clear from our analyses alone and may be due partly to other complex effects such the suppression of herbivory from sediments [46]. The results also indicate some negative effects of land-based pollution on benthic condition (Fig 3a), although we found mixed results for the direct effects of wastewater effluent and sediment that were driven by single anomalous locations, complicating our interpretation of these sources land-based pollution on the benthos. These patterns reflect the difficulty in aligning spatial and temporal scales of chronic drivers and spatial benthic patterns [47]. Nonetheless, habitat restoration, minimizing urban runoff, and reducing cesspool effluent to decrease the negative effects of land-based pollution may contribute to multiple goals of resilience-based management including direct improvements to benthic and herbivore assemblages, and indirect improvements for both due to the reciprocal relationships between them. Efforts focused on land-based pollution are thus all worthwhile endeavors; however, the mixed results for the direction of the effects of land-based pollution on benthic condition and the relatively stronger effects of herbivores on benthic condition also underscores the complexity of land-sea linkages [48]. It is likely necessary to consider where land and sea impacts are overlapping, and understand the tradeoffs and potential benefits associated with coordinated actions on land and at sea [49], especially in a changing climate [50].

### Conclusion

By combining reef survey data from multiple agencies and institutions with comprehensive driver data, this research highlights spatial patterns in herbivores and benthic condition for coral reefs across Hawai’i at spatial scales relevant for management. Combining spatiotemporal associations with the analysis of management interventions allows this study to inform ongoing management initiatives addressing both fishing and land-based drivers of coral reef resilience. Our results highlight the association between fishing pressure and herbivorous fish biomass that influence reef condition, and present evidence for fisheries interventions that increase fish biomass. These links between herbivores, fisheries management, and benthic condition pre-date substantial climate driven heat stress events that began in 2014 in Hawai’i, so further work is needed to understand whether the fishing threshold found here translates to reef resilience to climate change. As management actions to address both fishing pressure and land-based pollutants are implemented, continued collaborative monitoring across agencies will inform and, thereby, empower cooperative management strategies for both local communities and the coral reef ecosystem.

## Methods

### (a) Ecological Data Compilation

Data on herbivore biomass and benthic condition were compiled from existing underwater visual reef census datasets collected by researchers at governmental, non-governmental, and academic institutions (Table S2) [28,36]. Fish counts were calibrated using species and method specific adjustments following Friedlander et al. [28] to account for differences among institutions in how researchers surveyed reef fish. We calculated fish biomass using the allometric equation: W=*a*TL^*b*^, where *a* and *b* are species-specific parameters obtained from Hawai’i data (unpublished) or Fishbase [51], W is weight in grams, and TL is total length in centimeters. When calculating survey level biomass, observations of schooling species that can lead to spurious biomass predictions were adjusted by calculating the upper 99.9% of all individual observations, resulting in 26 observations out of over 0.5 million, comprised of 11 species (including 7 herbivore species). We then adjusted the counts for those 7 species where the individual count fell about the 99% quantile across all observations by truncating the count to the 99% quantile.

We subset the combined surveys to those focused on reef habitats, which we have defined as hard-bottom habitats from 0-30 meters depth, in main Hawaiian islands. Data were further subset to those collected between 2004 and 2014 to capture the condition of Hawai’i reefs prior to a major coral bleaching event in 2014-15 that caused widespread coral declines [26]. Thus, our analyses are for the condition of reefs prior to this pulse event and allow for inferences to be made about reefs that forestall a ‘shifting baseline’ where already-degraded reefs become normative [52]. The 2004-2014 period included 20,242 unique fish transects that we aggregated into 5,770 ‘replicates’, defined as data collected by a given data source at a unique latitude, longitude, depth, and year (Table S2) with means across transects taken where multiple transects occurred within each replicate. Of those, 1,182 (20% of the data) replicates were located within 2.5 km of each other at Kahekili reef, a marine managed area that has been extensively monitored [53]. We conducted sensitivity analyses to assess the potential for this high density of points in one area to affect our inferences (Fig S6), and subsequently used only 2014 data from Kahekili (n=159) in models of benthic condition and herbivore assemblages in Figures 1-3. Of the 4,775 replicates 2,029 had coincident benthic data collected that were used in analyses of herbivore-benthic relationships (Fig 3, Table S2).

### (b) Metrics of herbivore assemblages and benthic condition

We calculated four herbivorous fish population metrics including total herbivore biomass and biomass of three key herbivore functional groups: grazers, scrapers, and browsers. Grazers (e.g., surgeonfishes) feed on algal turf and small fleshy algae, and scraper/excavators (e.g., parrotfishes) on underlying substrate, thus helping to reduce colonization by macroalgae [25]. Therefore, a high biomass of grazers and scrapers may be particularly important for maintaining the reef in a calcified state. Browsers (e.g., chubs) are also important because they feed primarily on mature macroalgae and may reverse an algal-dominated state [54]. Browsers may play an important role in the recovery of coral reefs, and a healthy proportion of browsers is necessary to maintain the reef in a calcified state for the long term. Fish species were assigned to herbivorous functional groups following Donovan et al. [36] (Table S3).

We characterized benthic condition using the log ratio of calcified to macroalgal cover. Coral cover is a widely used indicator that captures overall patterns in the prevalence of corals, which are keystone species and primary habitat builders on coral reefs. However, coral cover is limited as a benthic indicator in that it does not differentiate pattern from process [55], so it is critical to capture other metrics of benthic condition including macroalgal cover and the cover of crustose coralline algae. Given this, we calculated the log-ratio of calcified to macroalgal cover as an integrated measure of reef benthic condition. Similar metrics have been shown to be an effective indicator of reef condition across human impact gradients [56]. Calcified cover is the sum of coral and crustose coralline algae cover, and we chose contrast calcified cover to macroalgal cover rather than ‘fleshy’ cover given that turf algae are ubiquitous on Hawai’i reefs [57] and our data does not differentiate between types of turf, which can vary greatly in functional roles [58,59]. A log transformation of the ratio was used to achieve parallel distributions around zero, allowing for the interpretation of sites dominated by calcifying organisms to have positive values and sites dominated by macroalgae to have negative values.

### (c) Drivers of herbivore assemblages

To examine the influence of human and biophysical factors on herbivore assemblage metrics, we compiled spatially comprehensive layers of potential predictor variables (Table S4). For metrics of land-based pollution (e.g., runoff, sedimentation), commercial and non-commercial fishing (e.g., boat-based net, shore-based spear), and physical oceanography (e.g., sea surface temperature, irradiance), we relied on previously created maps from Lecky [60] and Wedding & Lecky et al. [61]. To consider variation in habitat types, we used vector-based habitat maps produced by NOAA’s Biogeography Branch [62] and considered four categories: reef (coral-dominated hardbottom), pavement, boulder, and other hardbottom. We also included depth and rugosity as important determinates of habitat features; however, depth and rugosity data were not available in a spatially comprehensive way for the entire study domain, so we synthesized multiple sources of data (Supplemental Methods).

For some of the predictor variables, multiple metrics (e.g., mean, max, standard deviation) were previously calculated and available. To avoid overfitting our model while retaining unique predictor variables that account for distinct processes, we grouped all predictor variables into 1) land-based pollution, 2) fishing, 3) physical oceanography, and 4) habitat factors. We then investigated correlations across all variables with particular attention paid to highly correlated variables within each group. For any variables that were correlated above Spearman’s ρ of 0.7, we selected one variable to retain in the model. This decision was based on which variable more closely represented our best hypothesis for an effect on herbivore biomass and which variable had a lower correlation to remaining variables. No variables across the four driver groups were correlated above Spearman’s ρ of 0.7. Our final predictor set included 27 variables (Table S4), which were processed to a consistent 100 m grid (Supplemental Methods).

### (d) Herbivore-driver relationships

For each of the four metrics of herbivore biomass, we created hierarchical Bayesian models that were a function of human and environmental variables while accounting for variation in space, time, and data source.

We parameterized the biomass models in terms of a Gamma distribution with the mean (μ) using 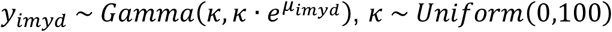. We then modeled the mean as a linear function of the multiple predictors (*β*) described in Table S4. All continuous predictors were standardized to a zero mean and unit variance to improve model convergence across predictors with different units. We included hierarchical effects of year (*y*) to account for variation over time, of moku (land-divisions, Fig S5) (*m*) to account for spatial variation, and of dataset (*d*) to account for additional effects of combining data from multiple methods and survey designs not captured by the calibration described previously, as:

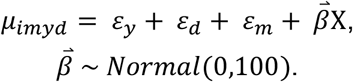

To account for low sampling across some moku, we removed data from moku with less than five replicates, bringing the final sample size used to fit the biomass models to 4,775 replicates. In order to ensure identifiability of the hierarchical effects, we implemented a ‘sum-to-zero-constraint’ as described by Ogle & Barber [63]. In summary, when hierarchical effects variance is large relative to the sample variance, this can create ‘nearly’ non-identifiable parameters as a result of implementation of Markov chain Monte Carlo (MCMC) methods that result in correlated posteriors. To address this, we constrained the average of the hierarchical effect to zero. For example, the hierarchical effect of year was:

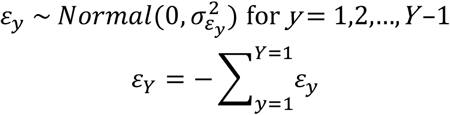

The models were fit with JAGS via the *rjags* package in R (Plummer 2016) with 7,500 iterations, including an initialization of 500 iterations, a burn-in of 2,000 iterations, with posterior estimates based on the remaining 5,000 iterations. Model convergence was assessed by running three chains and calculating Gelman-Rubin statistics [64] (Table S5). Model fits were assessed with posterior predictive checks following [65] and Bayesian R^2^ (Fig S7).

### (e) Spatial variation of herbivore assemblages

Model inputs were based on geographic coordinates where survey data were located. Because the multi-institution reef surveys were designed for many different purposes, the resulting combined dataset is not evenly distributed across space and, therefore, is not a balanced representation of habitats across the study domain. To produce appropriately-weighted herbivore biomass estimates, we post-stratified the predictions. That is, we predicted herbivore biomass based on drivers at a resolution of 100 m. This gives every prediction in the 100-m grid equal weight when summarized to larger areas (Fig 2, Fig S2).

To provide information at other spatial scales relevant to management (e.g., moku, management areas), we also summarized the spatial predictions to create post-stratified estimates that incorporate underlying spatial and model variability within each area of interest. To do so, we combined the full posterior samples for each 100 m^2^ pixel within each area of interest and calculated means and intervals across the combined posteriors. We removed pixels that were dominated by soft bottom (Supplemental Methods) from these estimates given that our sampling domain was restricted to hard bottom areas.

### (f) Benthic-driver relationships

To investigate drivers of benthic condition we constructed a model similar to the herbivore-driver model with the log ratio of calcified to macroalgal cover as the response and the biomass of the three herbivore functional groups and their interactions as drivers, along with the same oceanographic, land-based pollution, depth, and rugosity drivers used in the herbivore model. We did not include the fishing drivers in this model because we hypothesized that fishing would affect the benthos indirectly through herbivores, and we did not include the habitat categories since those are circular with the response variable. The benthic model was based on a subset of the data used previously where both fish and benthic data were collected on the same surveys, resulting in 2029 replicates (Table S3). Biomass of herbivore functional groups was Log+1 transformed to reduce the influence of extreme values and to improve model convergence.

We parameterized the model in terms of a Normal distribution with *y*_*imyd*_ ∼ *Normal*(μ, τ), τ ∼ *Gamma*(0.1,0.1) where the standard deviation *σ* is 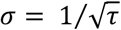 We then modeled the mean as a linear function of the transformed biomass of the three herbivore functional groups and their two- and three-way interactions and the other drivers. As previously, we included hierarchical effects of year (*y*) to account for variation over time, of moku (land-divisions) (*m*) to account for spatial variation, and of dataset (*d*) to account for additional effects of combining data from multiple methods and survey designs, using the same sum-to-zero constraint described previously. The model was fit with JAGS via the *rjags* package in R [66] with 7,500 iterations, including an initialization of 500 iterations, a burn-in of 2,000 iterations, with posterior estimates based on the remaining 5,000 iterations. Model convergence was assessed by running three chains and calculating Gelman-Rubin statistics [64] (Table S5). Model fits were assessed with posterior predictive checks (Fig S8) following [65] and Bayesian R^2^.

### (g) Patterns of fishing effects on herbivores and benthos

To understand how the magnitude and direction of fishing drivers of herbivores were related to benthic condition, we combined predictions from the herbivore-driver models and the benthic-driver model (Fig S9).

From the herbivore-driver model, for every 100 m^2^ pixel we calculated a posterior estimate of the ‘expected’ herbivore biomass for all four herbivore metrics from the measured drivers for that pixel. We also calculated a posterior estimate of the ‘potential’ biomass for each pixel in the absence of fishing by setting fishing drivers to zero (prior to scaling) and holding all other drivers at measured values. Finally, we calculated the “% of potential biomass” by dividing the posteriors of expected biomass by those of potential biomass for each pixel.

To understand the sensitivity of benthic condition to differences in herbivore functional group biomass from fishing we calculated a posterior estimate of the log-ratio of calcified to macroalgal cover for each pixel from the benthic model using the measured drivers and the mean of the posteriors for the three herbivore functional groups for both expected and potential biomass. We then classified each pixel as affected or unaffected by fishing depending on whether the 25% percentile of posterior log-ratio of calcified to macroalgal cover given potential biomass was greater than the 75% percentile given expected biomass (in other words, did the 50% interval overlap and was the interval for potential biomass greater than expected biomass). This binomial variable was then modeled as a function of the median posterior % potential biomass with a logistic regression to quantify the % potential biomass associated with a 99% probability that benthic condition is reduced from depletion of herbivores by fishing. We then compared this threshold value of % potential biomass to the distribution of values across the study domain and within and across moku. We chose the 99% probability to calculate the threshold given that the sample size was large (130,149 pixels) so we can rely on a relatively certain cutoff for determining what level of fishing might translate to decreased benthic condition.

### (h) Management effectiveness case studies

To understand how different management actions can influence herbivore populations, we analyzed patterns of parrotfish biomass in four management interventions (See Fig S5 for locations of survey sites). We focus on parrotfishes as they are heavily targeted by both commercial and non-commercial fisheries in Hawai’i [67], and they provide important ecological functions that can be compromised by overexploitation [45]. We tested whether herbivore populations increased following each intervention by comparing the biomass of parrotfishes using a binomial-Gamma hurdle model. Zero and non-zero data were both modeled as a function of management (e.g., before closure/after closure) and fit with the ‘glm’ function in R using either a binomial distribution with a logit link, or a Gamma distribution with a log link. The combined predictions with bias-corrected 95% confidence intervals were then calculated using non-parametric bootstrapping of the model 5,000 times with the ‘boot’ and ‘boot.ci’ functions in R.

Maui has several no-take marine reserves and island-wide rules implemented in 2014 specific to parrotfishes ([HAR 13.95.1](https://dlnr.hawaii.gov/dar/files/2014/10/ch95-1.pdf)), including prohibited take of terminal-phase males, size limits, and a bag-limit of two parrotfish per day per person, which reduced overall take of some species by up to 86% (State of Hawaii 2022). Data collected between 2018 and 2019 by the Hawai’i Division of Aquatic Resources at three marine reserves, and eight non-reserve sites, were used to test the effectiveness of these reserves using the binomial-Gamma hurdle model with management (open, protected) as a predictor.

The Kahekili Herbivore Fisheries Management Area, located on the western coastline of Maui in the Kā’anapali area (Fig S5), was established in 2009 in response to increasing algal cover and decreasing coral cover ([HAR 13-60.7](https://dlnr.hawaii.gov/dar/files/2014/05/ch60.7.pdf)). Take of herbivorous fishes and urchins is prohibited, but all other fishing is permitted. Kahekili was monitored extensively through a joint-agency effort from 2008-2016 [53]. We compared parrotfish biomass using the binomial-Gamma hurdle model between the years surveyed within the managed area before the closure (2008-2009) to the years surveyed following the closure (2011-2016).

The State of Hawai’i amended the rules of the West Hawai’i Regional Fishery Management Area (WHRFMA) in December 2013 to include a ban on SCUBA spearfishing anywhere within the WHRFMA boundaries (Fig S5), which extends along the entirety of Hawai’i Island’s western coast. Surveys of reef fishes at 25 permanent monitoring sites generally occurred four times a year within the WHRFMA [69]. We tested patterns of parrotfish biomass in the WHRFMA following the implementation of the SCUBA spearfishing ban at 10 sites not in protected areas, 13 sites inside Fishery Management Areas (aquarium collecting prohibited and other gear restrictions in some areas), and two sites in Marine Life Conservation Districts (no-take areas). Given the extensive time series, we compared four years of data 2007-2010 as ‘before’ the spearfishing ban and four years of data from 2016-2019 as ‘after’ the ban to provide a buffer of three years before and after for comparison.

## Data Availability Statement

All statistical analyses were performed in the R language for statistical computing version 4.0 [70]. Cartographic visualizations were made in ArcGIS Pro, and bathymetric lidar point cloud data processed with LAS Tools (https://rapidlasso.com/lastools/) and ArcGIS Desktop. All data and R scripts used to perform analyses and prepare figures can be accessed at https://doi.org/10.5281/zenodo.7478759 [71].

## Acknowledgements

We are grateful to the many divers who have surveyed Hawai’i reefs over the decade of data included in this study, and to the institutions that support nearshore monitoring that provide fish and benthic data to the Hawai’i Monitoring and Reporting Collaborative (HIMARC): Coral Reef Assessment and Monitoring Program (CRAMP) at the University of Hawai’i, National Oceanic and Atmospheric Administration (NOAA) US National Coral Reef Monitoring Program, State of Hawai’i Division of Aquatic Resources, University of Hawai’i Fisheries Ecology Research Laboratory including data from NOAA’s Fish Habitat Utilization Study, US National Park Service, and The Nature Conservancy Hawai’i. We are especially grateful to Eric Brown, Eric Co, Eric Conklin, Alan Friedlander, Jamison Gove, Mike Lameier, Brian Neilson, Ku’ulei Rodgers, and Ivor Williams, who have been steadfast supporters of HIMARC over the past decade, as well as our colleagues from the Ocean Tipping Points Project whose previous efforts contributed to the driver data. Rachel Layko was instrumental in wrangling data on soft bottoms to incorporate in our spatial predictions, and Ellie Jones assisted with Fig S9. We acknowledge Research Computing at Arizona State University for providing high-performance computing resources that have contributed to this research.

## Literature cited

1. van Hooidonk R, Maynard J, Tamelander J, Gove J, Ahmadia G, Raymundo L, Williams G, Heron SF, Planes S. 2016 Local-scale projections of coral reef futures and implications of the Paris Agreement. Sci. Rep. 6, 39666.

2. Halpern BS et al. 2008 A global map of human impact on marine ecosystems. Science 319, 948–952.

3. Fabricius KE. 2005 Effects of terrestrial runoff on the ecology of corals and coral reefs: review and synthesis. Mar. Pollut. Bull. 50, 125–146.

4. Jackson JBC, Donovan MK, Cramer KL, Lam V V. 2014 Status and Trends of Caribbean Coral Reefs: 1970-2012. IUCN, Gland, Switzerland.

5. Zaneveld JR et al. 2016 Overfishing and nutrient pollution interact with temperature to disrupt coral reefs down to microbial scales. Nat. Commun. 7, 11833.

6. Donovan MK, Burkepile DE, Kratochwill C, Shlesinger T, Sully S, Oliver TA, Hodgson G, Freiwald J, van Woesik R. 2021 Local conditions magnify coral loss after marine heatwaves. Science 372, 977–980.

7. Graham NAJ, Bellwood DR, Cinner JE, Hughes TP, Norström A V., Nyström M. 2013 Managing resilience to reverse phase shifts in coral reefs. Front. Ecol. Environ. 11, 541–548.

8. Anthony K et al. 2015 Operationalizing resilience for adaptive coral reef management under global environmental change. Glob. Chang. Biol. 21, 48–61.

9. Mcleod E et al. 2019 The future of resilience-based management in coral reef ecosystems. J. Environ. Manage. 233, 291–301.

10. Chung AE, Wedding LM, Meadows A, Moritsch MM, Donovan MK, Gove J, Hunter C. 2019 Prioritizing reef resilience through spatial planning following a mass coral bleaching event. Coral Reefs 38.

11. Chung A, Oliver T, Gove J, Gorospe K, White D, Davidson K, Walsh W. 2019 Translating resilience-based management theory to practice for coral bleaching recovery in Hawai’i. Mar. Policy 99, 58–68.

12. Mumby PJ, Wolff NH, Bozec YM, Chollett I, Halloran P. 2014 Operationalizing the resilience of coral reefs in an era of climate change. Conserv. Lett. 7, 176–187.

13. Steneck RS, Mumby PJ, MacDonald C, Rasher DB, Stoyle G. 2018 Attenuating effects of ecosystem management on coral reefs. Sci. Adv. 4, eaao5493.

14. Bellwood D, Hughes T, Folke C, Nyström M. 2004 Confronting the coral reef crisis. Nature 429, 827–833.

15. Burkepile DE, Hay ME. 2008 Herbivore species richness and feeding complementarity affect community structure and function on a coral reef. Proc. Natl. Acad. Sci. 105, 16201–16206.

16. Adam TC, Burkepile DE, Ruttenberg BI, Paddack MJ. 2015 Herbivory and the resilience of Caribbean coral reefs: knowledge gaps and implications for management. Mar. Ecol. Prog. Ser. 520, 1–20.

17. Williams ID, Kindinger TL, Couch CS, Walsh WJ, Minton D, Oliver TA. 2019 Can Herbivore Management Increase the Persistence of Indo-Pacific Coral Reefs? Front. Mar. Sci. 6, 557.

18. Cinner JE et al. 2020 Meeting fisheries, ecosystem function, and biodiversity goals in a human-dominated world. Science 368, 307–311.

19. Hughes TP, Rodrigues MJ, Bellwood DR. 2007 Phase shifts, herbivory, and the resilience of coral reefs to climate change. Curr. Biol. 17, 360–365.

20. Rasher DB, Engel S, Bonito V, Fraser GJ, Montoya JP, Hay ME. 2012 Effects of herbivory, nutrients, and reef protection on algal proliferation and coral growth on a tropical reef. Oecologia 169, 187–198.

21. Stockwell B, Jadloc CRL, Abesamis RA, Alcala AC, Russ GR. 2009 Trophic and benthic responses to no-take marine reserve protection in the Philippines. Mar. Ecol. Prog. Ser. 389, 1–15.

22. Wismer S, Hoey A, Bellwood DR. 2009 Cross-shelf benthic community structure on the Great Barrier Reef: relationships between macroalgal cover and herbivore biomass. Mar. Ecol. Prog. Ser. 376, 45–54.

23. Heenan A, Williams ID. 2013 Monitoring herbivorous fishes as indicators of coral reef resilience in American Samoa. PLoS One 8, e79604.

24. Roff G, Mumby PJ. 2012 Global disparity in the resilience of coral reefs. Trends Ecol. Evol. 27, 404–413.

25. Green AL, Bellwood DR. 2009 Monitoring functional groups of herbivorous reef fishes as indicators of coral reef resilience. IUCN Working Group on Climate Change and Coral Reefs, IUCN, Gland, Switzerland.

26. Kramer K, Cotton S, Lamson M, Walsh W. 2016 Bleaching and catastrophic mortality of reef-building corals along west Hawai ‘i island: findings and future directions. In Proceedings of the 13th International Coral Reef Symposium, Honolulu, pp. 229–241.

27. Friedlander AM, DeMartini EE. 2002 Contrasts in density, size, and biomass of reef fishes between the northwestern and the main Hawaiian islands: the effects of fishing down apex predators. Mar. Ecol. Prog. Ser. 230, 253–264.

28. Friedlander AM et al. 2018 Human-induced gradients of reef fish declines in the Hawaiian Archipelago viewed through the lens of traditional management boundaries. Aquat. Conserv. 28, 146–157.

29. Maire E et al. 2016 How accessible are coral reefs to people? A global assessment based on travel time. Ecol. Lett. 19, 351–360.

30. Cinner JE et al. 2018 Gravity of human impacts mediates coral reef conservation gains. Proc. Natl. Acad. Sci. 115, E6116–E6125.

31. Gove JM, Williams GJ, McManus MA, Heron SF, Sandin SA, Vetter OJ, Foley DG. 2013 Quantifying climatological ranges and anomalies for Pacific coral reef ecosystems. PLoS One 8.

32. Helyer J, Samhouri JF. 2017 Fishing and environmental influences on estimates of unfished herbivorous fish biomass across the Hawaiian Archipelago. Mar. Ecol. Prog. Ser. 575, 1–15.

33. Chong-Seng KM, Nash KL, Bellwood DR, Graham NAJ. 2014 Macroalgal herbivory on recovering versus degrading coral reefs. Coral Reefs 33, 409–419.

34. Randall JE. 1961 Overgrazing of algae by herbivorous marine fishes. Ecology 42, 812.

35. Jones RS. 1968 Ecological relationships in Hawaiian and Johnston Island Acanthuridae (surgeonfishes). Micronesica 4, 309–361.

36. Donovan MK et al. 2018 Combining fish and benthic communities into multiple regimes reveals complex reef dynamics. Sci. Rep. 8.

37. Russ GR, Questel S-LA, Rizzari JR, Alcala AC. 2015 The parrotfish–coral relationship: refuting the ubiquity of a prevailing paradigm. Mar. Biol. 162, 2029–2045.

38. Andradi-Brown DA et al. 2023 Diversity in marine protected area regulations: Protection approaches for locally appropriate marine management. Front. Mar. Sci. 10, 1099579.

39. Gaines SD, White C, Carr MH, Palumbi SR. 2010 Designing marine reserve networks for both conservation and fisheries management. Proc. Natl. Acad. Sci. 107, 18286–18293.

40. Spalding AK et al. 2023 Engaging the tropical majority to make ocean governance and science more equitable and effective. npj Ocean Sustain. 2, 8.

41. Williams ID, White DJ, Sparks RT, Lino KC, Zamzow JP, Kelly ELA, Ramey HL. 2016 Responses of Herbivorous Fishes and Benthos to 6 Years of Protection at the Kahekili Herbivore Fisheries Management Area, Maui. PLoS One 11, e0159100.

42. McClanahan TR, Graham NAJ, MacNeil MA, Muthiga NA, Cinner JE, Bruggemann JH, Wilson SK. 2011 Critical thresholds and tangible targets for ecosystem-based management of coral reef fisheries. Proc. Natl. Acad. Sci. U. S. A. 108, 17230–17233.

43. Karr KA, Fujita R, Halpern BS, Kappel C V, Crowder L, Selkoe KA, Alcolado PM, Rader D. 2015 Thresholds in Caribbean coral reefs: implications for ecosystem-based fishery management. J. Appl. Ecol. 52, 402–412.

44. Graham NAJ, Jennings S, MacNeil MA, Mouillot D, Wilson SK. 2015 Predicting climate-driven regime shifts versus rebound potential in coral reefs. Nature 518, 94–97.

45. Bozec Y-M, O’Farrell S, Bruggemann JH, Luckhurst BE, Mumby PJ. 2016 Tradeoffs between fisheries harvest and the resilience of coral reefs. Proc. Natl. Acad. Sci., 201601529.

46. Goatley CHR, Bellwood DR. 2012 Sediment suppresses herbivory across a coral reef depth gradient. Biol. Lett. 8, 1016–1018.

47. Donovan MK et al. 2023 From polyps to pixels: understanding coral reef resilience to local and global change across scales. Landsc. Ecol. 38, 737–752.

48. Nalley EM, Tuttle LJ, Barkman AL, Conklin EE, Wulstein DM, Richmond RH, Donahue MJ. 2021 Water quality thresholds for coastal contaminant impacts on corals: A systematic review and meta-analysis. Sci. Total Environ. 794, 148632.

49. Oleson KLL, Falinski KA, Lecky J, Rowe C, Kappel C V, Selkoe KA, White C. 2017 Upstream solutions to coral reef conservation: The payoffs of smart and cooperative decision-making. J. Environ. Manage. 191, 8–18.

50. Gove JM et al. 2023 Coral reefs benefit from reduced land–sea impacts under ocean warming. Nature, 1–7.

51. Froese R, Pauly D. 2011 Fishbase. http://www.fishbase.org.

52. Pauly D. 1995 Anecdotes and the shifting baseline syndrome of fisheries. Trends Ecol. Evol. 10, 430.

53. Williams ID, White DJ, Sparks RT, Lino KC, Zamzow JP, Kelly ELA, Ramey HL. 2016 Responses of Herbivorous Fishes and Benthos to 6 Years of Protection at the Kahekili Herbivore Fisheries Management Area, Maui. PLoS One 11, e0159100.

54. Bellwood DR, Hughes TP, Hoey AS. 2006 Sleeping functional group drives coral-reef recovery. Curr. Biol. 16, 2434–2439.

55. Bellwood DR et al. 2019 Coral reef conservation in the Anthropocene: confronting spatial mismatches and prioritizing functions. Biol. Conserv. 236, 604–615.

56. Smith JE et al. 2016 Re-evaluating the health of coral reef communities: baselines and evidence for human impacts across the central Pacific. Proc. R. Soc. B Biol. Sci. 283, 20151985.

57. Jouffray JB, Nyström M, Norström A, Williams ID, Wedding LM, Kittinger JN, Williams GJ. 2015 Identifying multiple coral reef regimes and their drivers across the Hawaiian archipelago. Philos. Trans. R. Soc. 370, 20130268.

58. Goatley CHR, Bonaldo RM, Fox RJ, Bellwood DR. 2016 Sediments and herbivory as sensitive indicators of coral reef degradation. Ecol. Soc. 21.

59. Bellwood DR, Streit RP, Brandl SJ, Tebbett SB. 2019 The meaning of the term ‘function’ in ecology: A coral reef perspective. Funct. Ecol. 33, 948–961.

60. Lecky J. 2016 Ecosystem vulnerability and mapping cumulative impacts on Hawaiian reefs. Thesis: University of Hawaii at Manoa.

61. Wedding LM et al. 2018 Advancing the integration of spatial data to map human and natural drivers on coral reefs. PLoS One 13.

62. Battista TA, Costa BM, Anderson SM. 2007 Shallow-water benthic habitats of the main eight Hawaiian Islands. NOAA Tech. Memo. NOS NCCOS 61.

63. Ogle K, Barber JJ. 2020 Ensuring identifiability in hierarchical mixed effects Bayesian models. Ecol. Appl. 30, e02159.

64. Gelman A, Rubin DB. 1992 Inference from iterative simulation using multiple sequences. Stat. Sci., 457–472.

65. Gabry J, Simpson D, Vehtari A, Betancourt M, Gelman A. 2019 Visualization in Bayesian workflow. J. R. Stat. Soc., A: Stat. Soc. 182, 389–402.

66. Plummer M. 2016 rjags: Bayesian Graphical Models using MCMC. R package version 4-6. https://CRAN.R-project.org/package=rjags.

67. McCoy KS, Williams ID, Friedlander AM, Ma H, Teneva L, Kittinger JN. 2018 Estimating nearshore coral reef-associated fisheries production from the main Hawaiian Islands. PLoS One 13, e0195840.

68. Resources HD of A. 2022 Rule Amendment Submittal HAR 13-95 Herbivore Rules.

69. Tissot BN, Walsh WJ, Hallacher LE. 2004 Evaluating effectiveness of a marine protected area network in West Hawai’i to increase productivity of an aquarium fishery. Pacific Sci. 58, 175–188.

70. R Core Team. 2019 R: A Language and Environment for Statistical Computing.

71. Donovan MK. 2023 Data and Code for Evidence for managing herbivores for reef resilience. Zenodo. (doi:10.5281/zenodo.7478759)

