## Supplemental Material for "Evidence for managing herbivores for reef resilience"

Contents:

Figures S1-S9
Tables S1-S5
Supplemental Methods

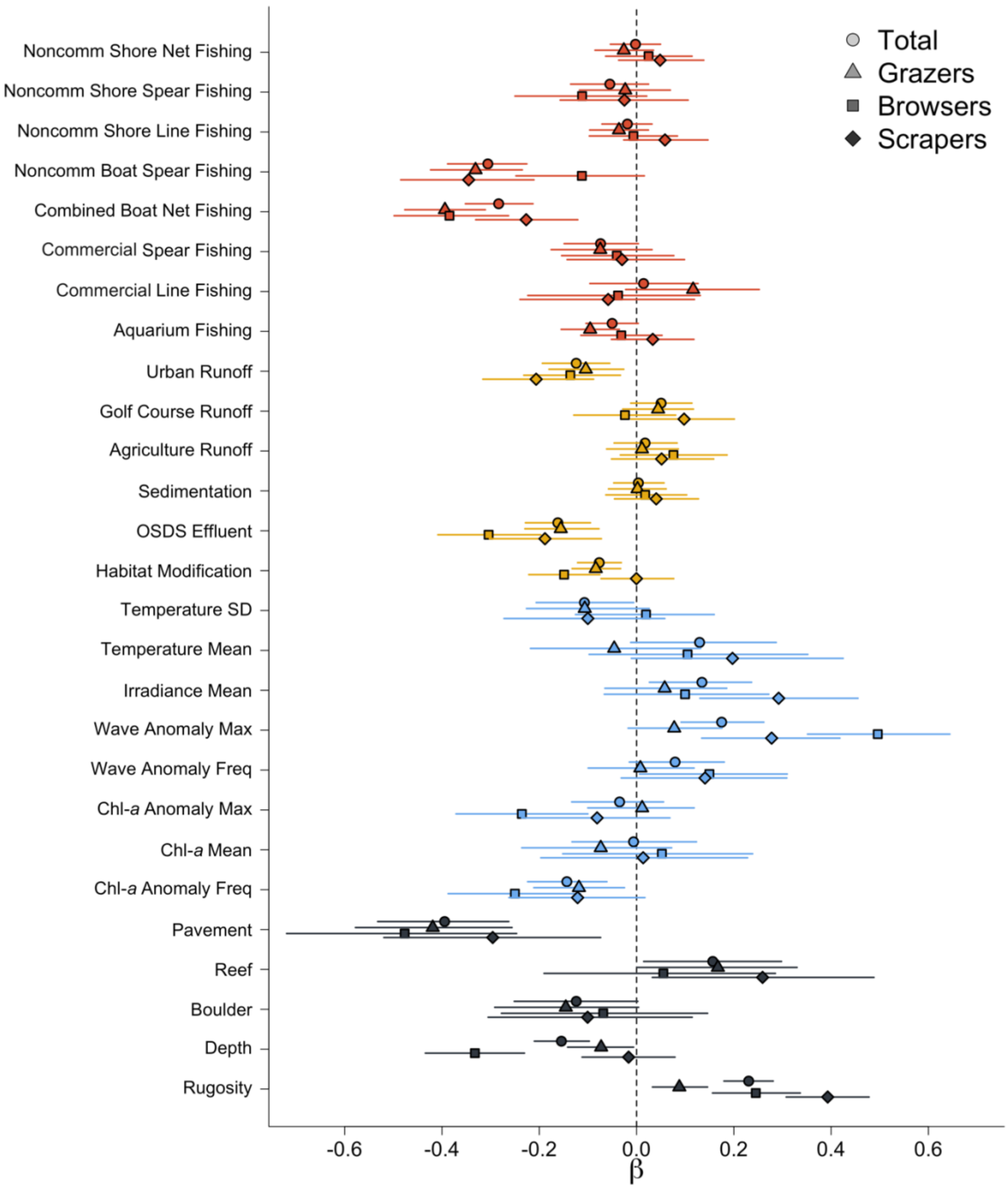


Figure S1. Coefficients ($\beta$) from Bayesian Hierarchical models estimating the relationship between drivers and total herbivore biomass (circles), grazer biomass (triangles), browser biomass (squares), and scraper biomass (diamonds). Points are median posterior estimates and horizontal lines are 95% intervals.

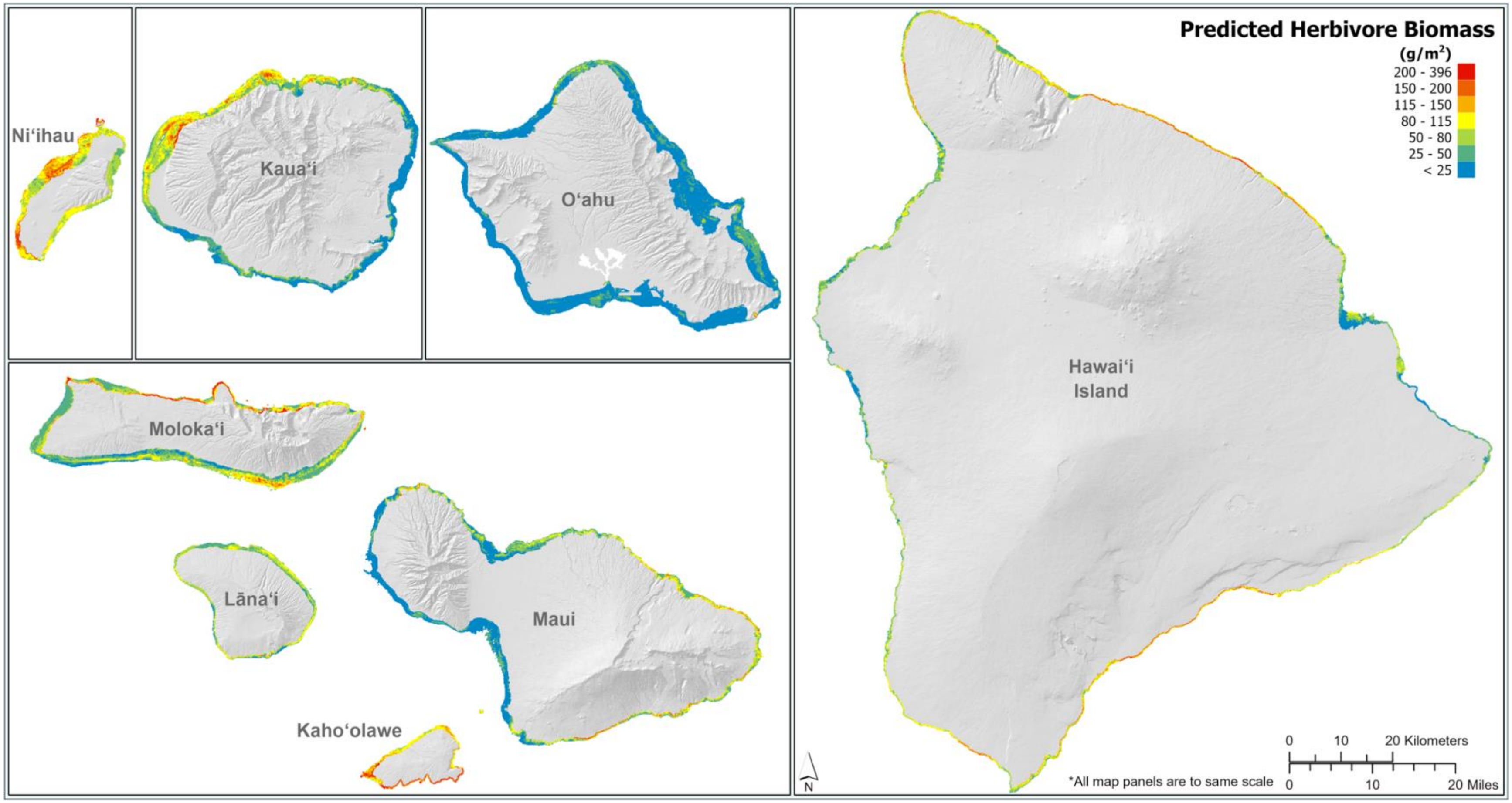


Figure S2. Predicted herbivore biomass across the study domain that includes nearshore waters from 0 to 30 m depth with the highest values in red and lowest values in blue. Predictions are based on driver values for each 100-m pixel and mean posterior estimates for all coefficients plus an offset based on the mean posterior estimate for each moku (land division).

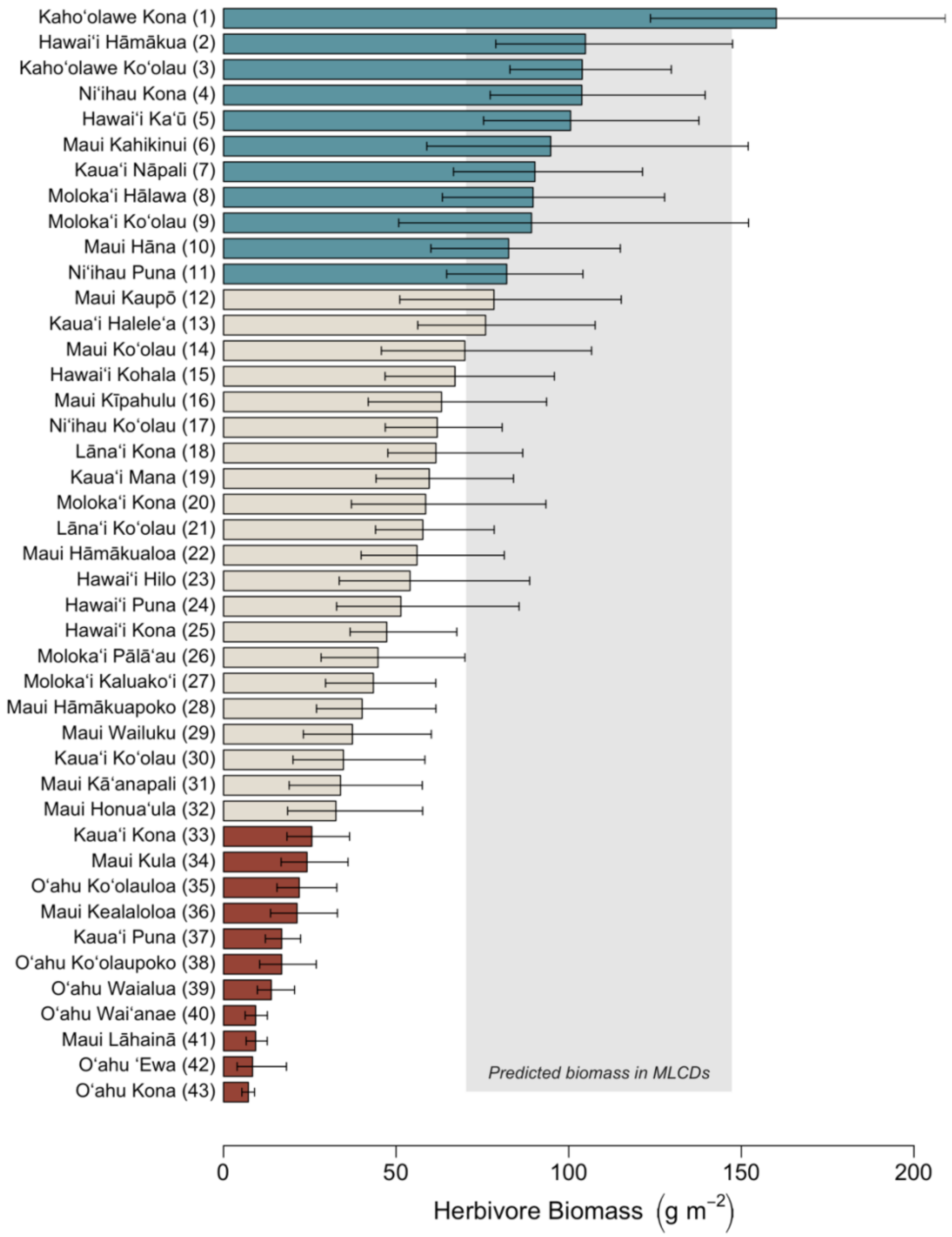


Figure S3. Posterior estimates of herbivore biomass summarized across moku (land divisions) (as plotted in Fig 2a). Bars are means and error bars are 50% intervals of posterior predictions for all 100 m pixels in each moku, and thus are post-stratified estimates that account for the relative distribution of habitat and variation in other predictors. Moku with mean values in the upper quartile (top fourth) are colored turquoise, and moku with mean values in the lower quartile (bottom forth) are colored red. Grey shaded area corresponds to the 50% interval of combined posteriors for all 100 m pixels inside Marine Life Conservation Districts, which prohibit fishing of herbivorous fishes.

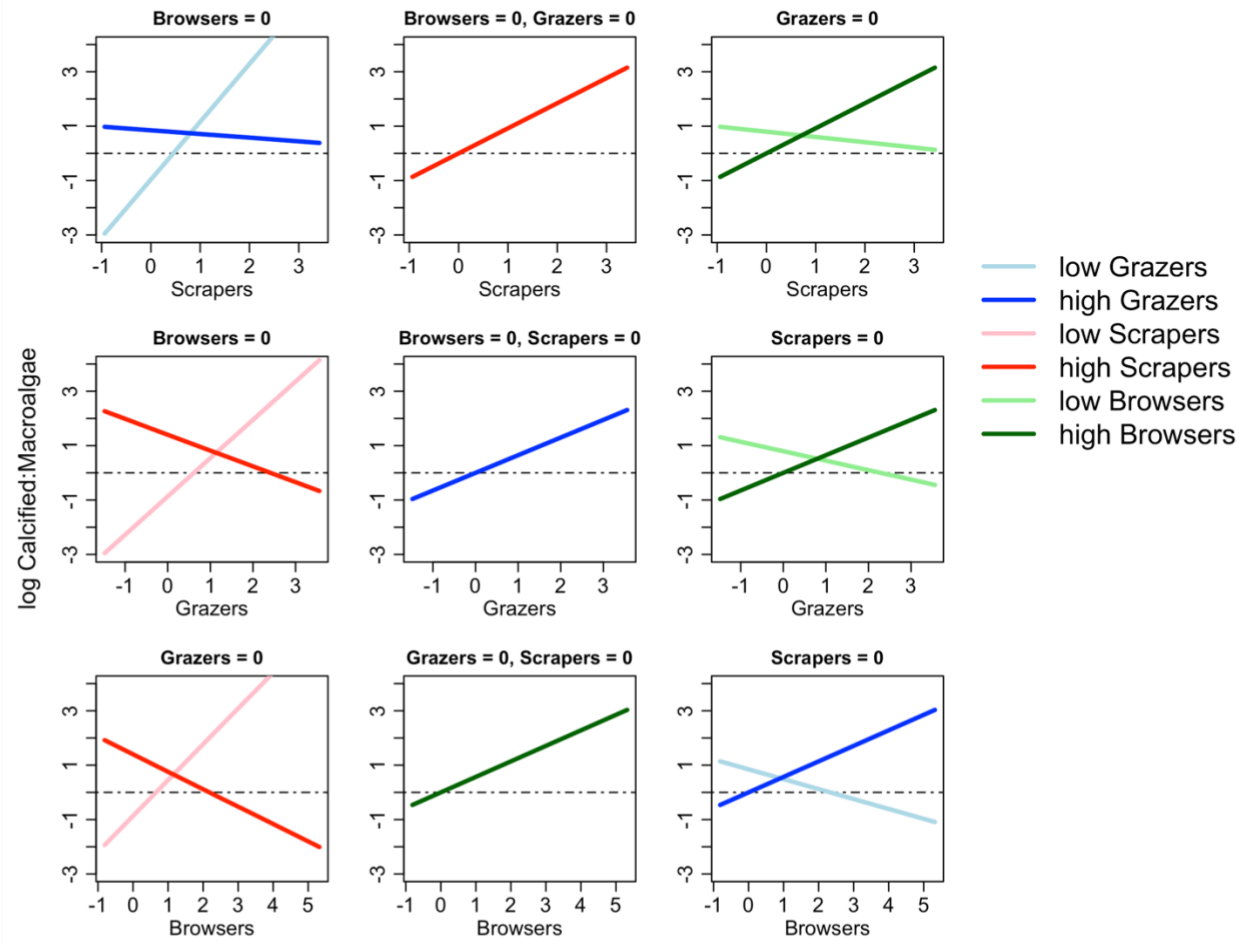


Figure S4. Interactions between herbivore functional group biomass as predictors of log-ratio of calcified and macroalgal cover. Values on the x-axis are herbivore functional group biomass log transformed and scaled reflecting the units as included in the model. Each panel displays the marginal effects with the values for each functional group held to either zero (panel label) or a high (75% quantile) or low (25% quantile) value (legend) with all other predictors held at their means.

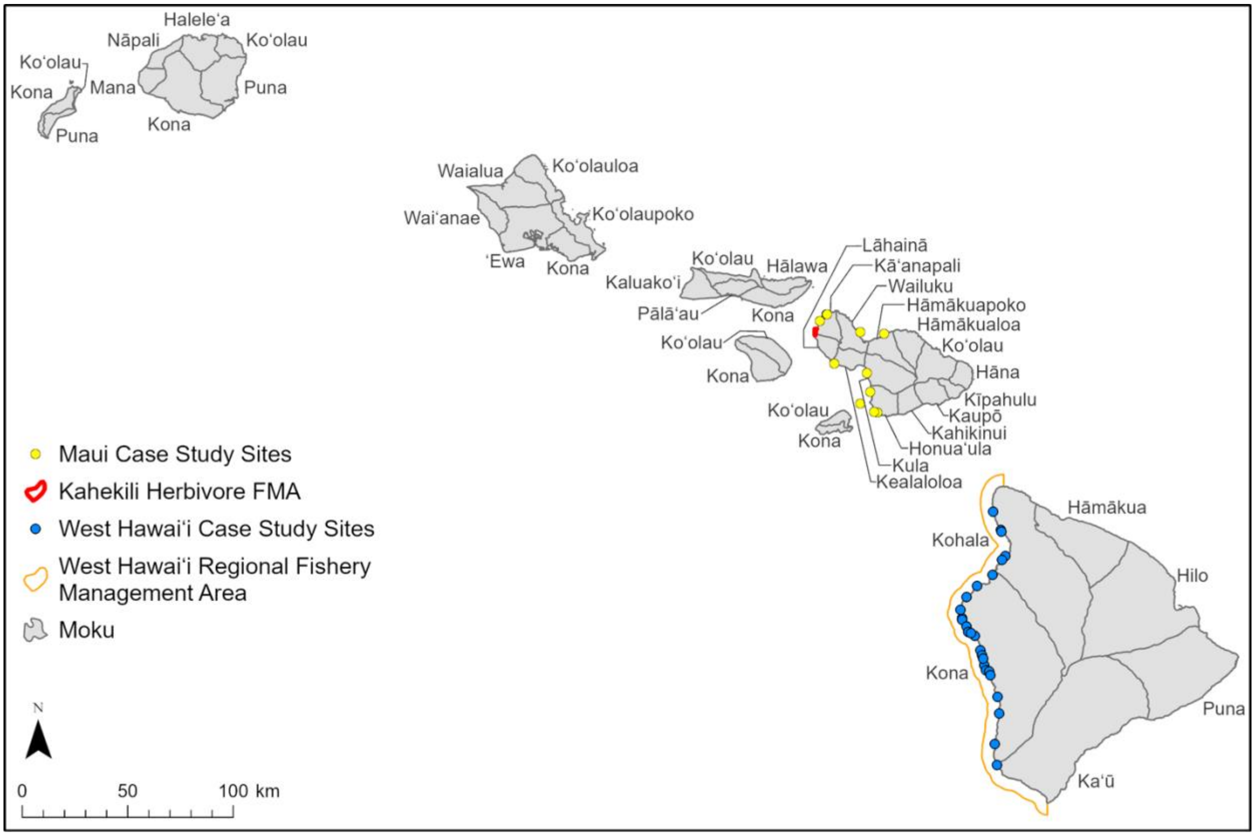


Figure S5. Map of the Main Hawaiian Islands with moku land divisions and survey site locations for data used in management effectiveness case study analyses.

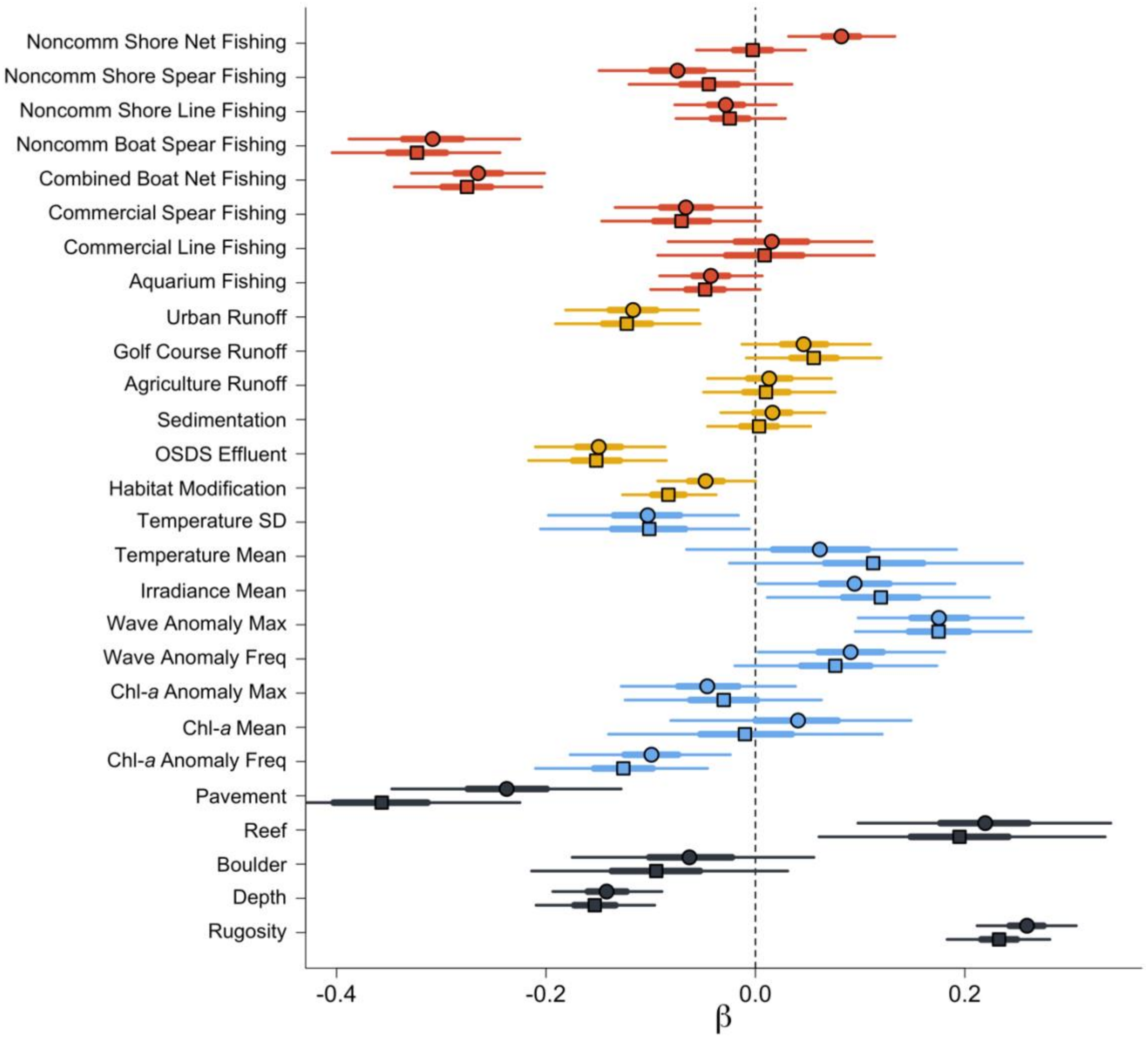


Figure S6. Comparison of results presented in Figure 1 across two subsets of data. Squares are the same results as Figure 1, with a subset of data excluded from the area of Kahekili reef in West Maui where intensive sampling has occurred accounting for 20% of the overall dataset. Circles are results from the same model run with all Kahekili data, showing that some relationships are driven by those data, particularly non-commercial shore net fishing.

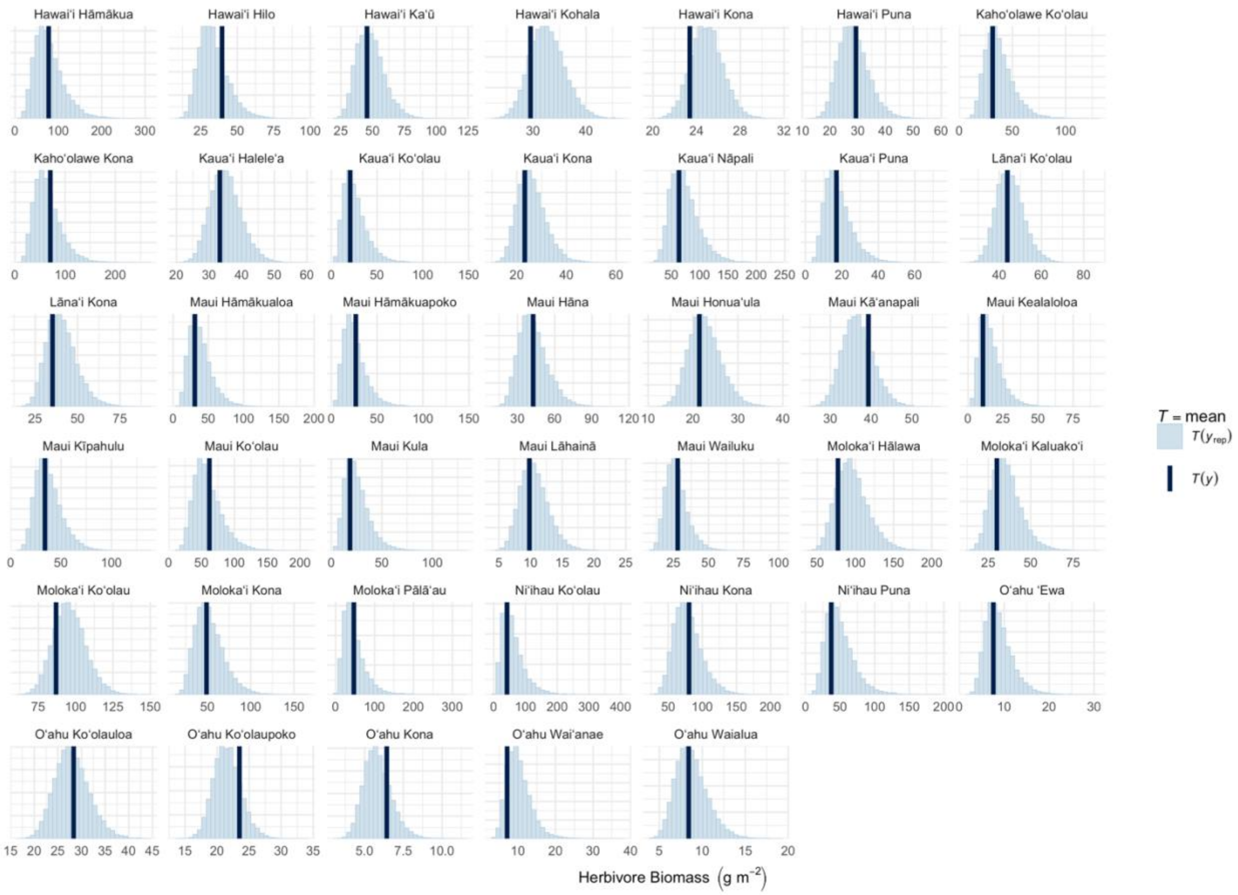


Figure S7. Posterior predictive check comparing empirical distributions of the mean of the observed data (vertical thick line) to distributions of the mean simulated from the posterior predictive distributions (light blue histograms) across moku (panels) for model of total herbivore biomass. The model is considered to reliably reproduce the test statistic (mean) when the vertical line is centered on the posterior predictive distributions. The x-axis is total herbivore biomass (g $m^{-2}$).

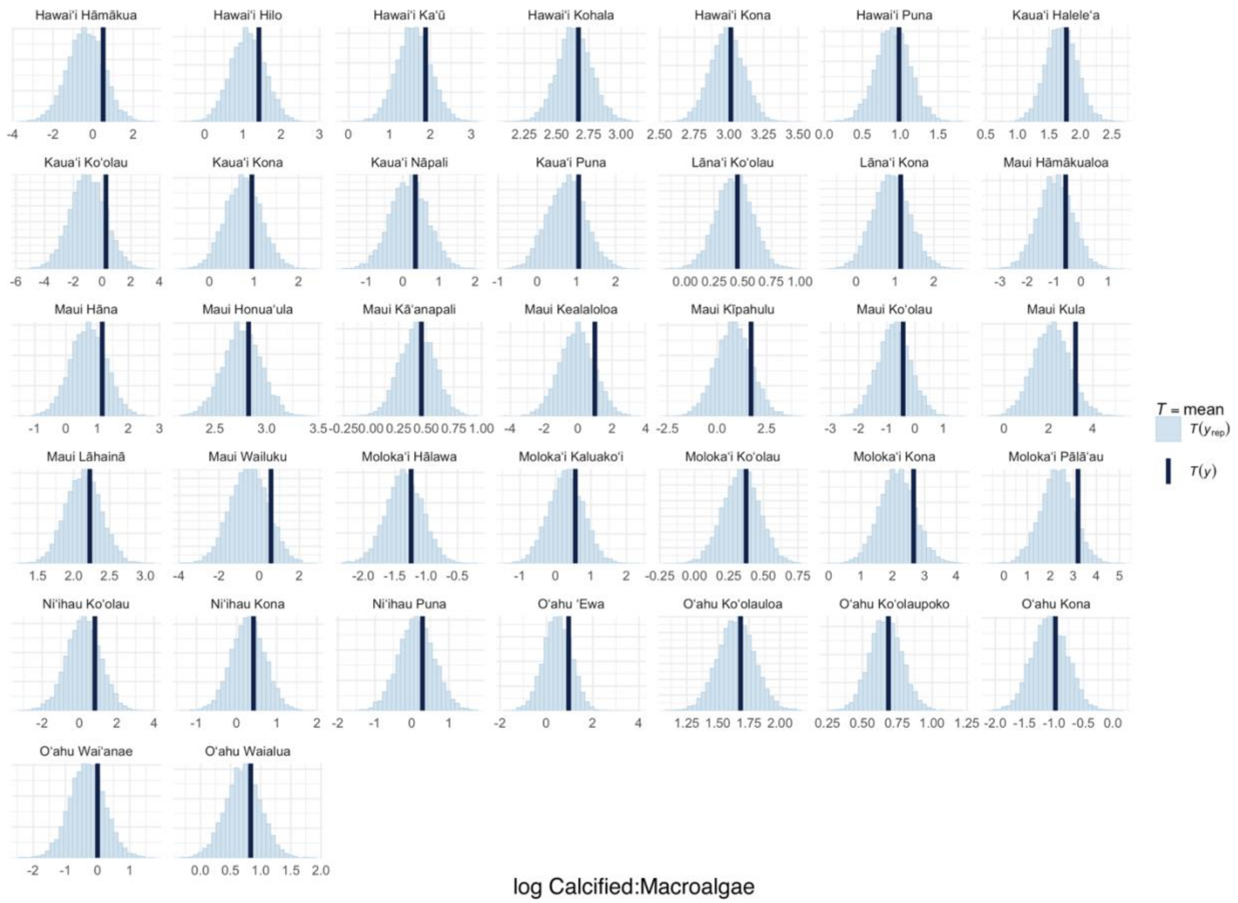


Figure S8. Posterior predictive check comparing empirical distributions of the mean of the observed data (vertical thick line) to distributions of the mean simulated from the posterior predictive distributions (light blue histograms) across moku (panels) for model of log-ratio of calcified to macroalgal cover. The model is considered to reliably reproduce the test statistic (mean) when the vertical line is centered on the posterior predictive distributions. The x-axis is log-ratio of calcified to macroalgal cover.

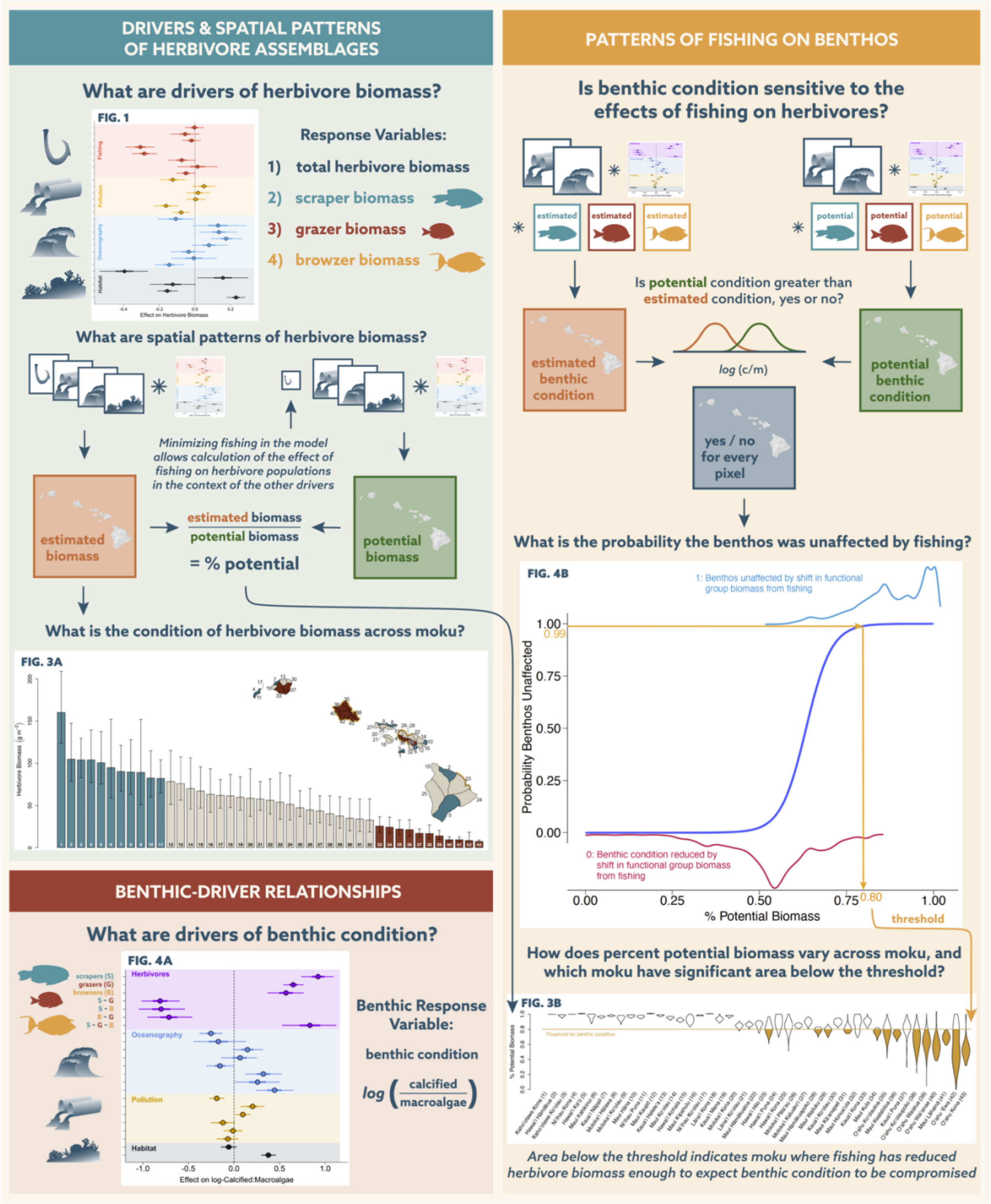


Figure S9. Infographic showing steps taken in analyses.

Table S1. Results of statistical models assessing management regulations.

**Maui, inside-outside**

| *Presence-absence* | Estimate | Std Error | P-value |
|---|---|---|---|
| Intercept | 0.11 | 0.07 | 0.135 |
| Management - Inside | 1.59 | 0.21 | <0.001 |
| *Biomass when present* | | | |
| Intercept | 2.73 | 0.07 | <0.001 |
| Management - Inside | 0.02 | 0.13 | 0.875 |

**Kahekili, before-after**

| *Presence-absence* | Estimate | Std Error | P-value |
|---|---|---|---|
| Intercept | -0.16 | 0.12 | 0.196 |
| Management - After | 1.04 | 0.14 | <0.001 |
| *Biomass when present* | | | |
| Intercept | 2.87 | 1.22 | <0.001 |
| Management - After | 0.27 | 0.13 | 0.039 |

**West Hawai‘i, non-reserve sites, before-after**

| *Presence-absence* | Estimate | Std Error | P-value |
|---|---|---|---|
| Intercept | 2.01 | 0.25 | <0.001 |
| Management - After | 0.87 | 0.44 | 0.046 |
| *Biomass when present* | | | |
| Intercept | 3.31 | 0.09 | <0.001 |
| Management - After | 0.21 | 0.12 | 0.089 |

**West Hawai‘i, fisheries management area sites, before-after**

| *Presence-absence* | Estimate | Std Error | P-value |
|---|---|---|---|
| Intercept | 2.63 | 0.28 | <0.001 |
| Management - After | 0.19 | 0.42 | 0.651 |
| *Biomass when present* | | | |
| Intercept | 3.43 | 0.07 | <0.001 |
| Management - After | 0.17 | 0.10 | 0.076 |

**West Hawai‘i, MLCD sites, before-after**

| *Biomass when present* | Estimate | Std Error | P-value |
|---|---|---|---|
| Intercept | 3.61 | 0.18 | <0.001 |
| Management - After | 0.68 | 0.25 | <0.001 |

Table S2. Data sources for herbivore biomass and benthic condition. Number of replicates is defined as data collected by a given data source at a unique latitude, longitude, depth, and year, with value in parentheses indicating the number of replicates at Kahekili reef.

| Name | Method(s) | Number Replicates | | Years | Islands |
|---|---|---|---|---|---|
| | | Fish | Benthic | | |
| Coral Reef Assessment and Monitoring Program (CRAMP) | Fish: 20 x 4 meter belt transects[1] | 48 | | 2012 | Moloka‘i, Kauai |
| National Oceanic and Atmospheric Administration US National Coral Reef Monitoring Program | Fish: 25 x 5 meter belt transect[2] or 15 meter diameter stationary point count[3]; Benthic: photo-quadrat transects[3] | 947 | 397 | 2005-2013 | Ni‘ihau, Kauai, O‘ahu, Moloka‘i, Lāna‘i, Maui, Hawai‘i |
| State of Hawai‘i Division of Aquatic Resources | Fish: 25 x 4 meter belt transects[4] or 25 x 5 meter belt transects[5]; Benthic: photo-quadrat transects[5] | 1707 (1023) | 813 | 2004-2014 | O‘ahu, Lāna‘i, Maui, Hawai‘i |
| University of Hawai‘i Fisheries Ecology Research Laboratory | Fish: 25 x 5 meter belt transects[5]; Benthic: in situ quadrat transects[5] | 548 | 290 | 2004-2014 | Kauai, O‘ahu, Lāna‘i |
| Fish Habitat Utilization Study | Fish: 25 x 5 meter belt transects[5]; Benthic: in situ quadrat transects[5] | 993 | 449 | 2004-2008 | O‘ahu, Maui, Hawai‘i |
| US National Park Service | Fish: 25 x 5 meter belt transects[5]; Benthic: photo-quadrat transects[6] | 548 | 120 | 2004-2014 | Moloka‘i, Hawai‘i |
| The Nature Conservancy Hawai‘i | Fish: 25 x 5 meter belt transects[5]; Benthic: photo-quadrat transects[7] | 979 | 187 | 2009-2014 | O‘ahu, Maui, Kaho‘olawe, Hawai‘i |

[1] Friedlander, A. M., Brown, E. E., Jokiel, P. I., Smith, W. R. & Rodgers, K. S. Effects of habitat, wave exposure, and marine protected area status on coral reef fish assemblages in the Hawaiian Archipelago. *Coral Reefs* **22,** 291–305 (2003).

[2] Williams, I. D., Walsh, W. J., Schroeder, R. E., Friedlander, A. M., Richards, B. L., & Stamoulis, K. A. (2008). Assessing the importance of fishing impacts on Hawaiian coral reef fish assemblages along regional-scale human population gradients. *Environmental Conservation*, *35*(3), 261-272.

[3] Williams, I. D. *et al.* Human, oceanographic and habitat drivers of Central and Western Pacific coral reef fish assemblages. *PLoS One* **10,** e0120516 (2015).

[4] Tissot, B. N., Walsh, W. J. & Hallacher, L. E. Evaluating effectiveness of a marine protected area network in West Hawai'i to increase productivity of an aquarium fishery. *Pacific Sci.* **58,** 175–188 (2004).

[5] Friedlander, A. M., Brown, E. & Monaco, M. E. Defining reef fish habitat utilization patterns in Hawaii: comparisons between marine protected areas and areas open to fishing. *Mar. Ecol. Prog. Ser.* **351,** 221–233 (2007).

[6] Brown, E. K. *et al.* Development of benthic sampling methods for the Coral Reef Assessment and Monitoring Program (CRAMP) in Hawai'i. *Pacific Sci.* **58,** 145–158 (2004).

[7] Stamoulis, K. A., & Friedlander, A. M. A seascape approach to investigating fish spillover across a marine protected area boundary in Hawai‘i. *Fisheries Research* **144**, 2-14 (2013).

Table S3. Species associated with each of the herbivore functional groups used in analyses.

| Browsers | Grazers | Scrapers |
|---|---|---|
| *Calotomus carolinus* | *Acanthurus achilles* | *Chlorurus perspicillatus* |
| *Calotomus zonarchus* | *Acanthurus blochii* | *Chlorurus spilurus* |
| *Kyphosus bigibbus* | *Acanthurus dussumieri* | *Scarus dubius* |
| *Kyphosus cinerascens* | *Acanthurus guttatus* | *Scarus psittacus* |
| *Kyphosus hawaiiensis* | *Acanthurus leucopareius* | *Scarus rubroviolaceus* |
| *Kyphosus vaigiensis* | *Acanthurus lineatus* | |
| *Naso lituratus* | *Acanthurus maculiceps* | |
| *Naso unicornis* | *Acanthurus nigricans* | |
| | *Acanthurus nigrofuscus* | |
| | *Acanthurus nigroris* | |
| | *Acanthurus olivaceus* | |
| | *Acanthurus triostegus* | |
| | *Acanthurus xanthopterus* | |
| | *Zebrasoma flavescens* | |
| | *Zebrasoma veliferum* | |

Table S4. Driver variables and data sources.

| Driver variable | Description | Data source |
|---|---|---|
| Land-Based Pollution | | |
| Urban runoff | Trash, household chemicals, oil from roads, and other forms of urban runoff were modeled as a proxy by calculating the area of impervious surfaces from the NOAA Coastal Change Analysis Program (CCAP) high resolution land use land cover (2010) per watershed and dispersing those values offshore. | Lecky 2016 |
| Golf course runoff | Pesticides and fertilizers from golf courses were modeled as a proxy using subsets from CCAP 'open developed space' land use class, and validated with Google Earth and ESRI Imagery, to calculate golf course area per watershed and disperse those values offshore. | Lecky 2016 |
| Agriculture runoff | Pesticides and fertilizers from agricultural runoff were modeled as a proxy by calculating the area of agricultural land from CCAP per watershed and dispersing those values offshore. | Lecky 2016 |
| Coastal-Habitat modification | Direct alteration, removal, and destruction of habitat including coastal engineering (e.g., seawalls, piers), dredging, and offshore aquaculture were compiled and represented as presence and absence. | Lecky 2016, Wedding et al. 2018 |
| Sedimentation | Modeled with the Integrated Valuation of Ecosystem Services and Tradeoffs (InVEST) sediment delivery ratio model to estimate the annual average delivery of sediment offshore. | Lecky 2016, Wedding et al. 2018 |
| On-site waste disposal effluent (OSDS) | Effluent from on-site waste disposal systems (cesspools, septic tanks) from estimated flux (gallons of wastewater per day) by land parcel from the Hawai‘i Department of Health and proximity to individual systems. | Lecky 2016, Wedding et al. 2018 |
| Fishing | | |
| Non-commercial boat-based spear | Island-scale annual average non-commercial reef fisheries catch (kg/ha) from 2004-2013 was calculated from the Marine Recreational Information Program data. Values were mapped offshore by modeling the distance to harbors and boat launches, human population within 30 km, and managed area regulations. | McCoy et al. 2018, Lecky 2016, Wedding et al. 2018 |
| Non-commercial shore-based spear, line, and net | Island-scale annual average non-commercial shore-based reef fisheries catch (kg/ha) by spear, line, and net from 2004-2013 were calculated from MRIP data. Values were mapped offshore by modeling shoreline accessibility using TIGER roads, USGS DEM slope, gear-specific spatial footprints, and managed area regulations. | Lecky 2016, Wedding et al. 2018 |
| Aquarium collection | Average annual reported commercial aquarium catch (#/ha) from 2003-2015 by reporting block from the Hawai‘i Division of Aquatic Resources. | Lecky 2016 |
| Commercial line<br>Commercial spear | Average annual commercial catch of reef fish species (kg/ha) with line and spear gear types by reporting block from 2003-2013 from Hawai‘i Division of Aquatic Resources. | Lecky 2016, Wedding et al. 2018 |

| | | |
|---|---|---|
| Boat-based net | Summed average annual catch of reef fish species by net from commercial reporting (as described above) and from non-commercial boat-based estimates (as described above). | Lecky 2016, Wedding et al. 2018 |
| Physical Oceanography | | |
| Temperature | Sea surface temperature from weekly 5 km NOAA blended satellite data from 2000-2013. Long-term mean and standard deviation of long-term mean were retained to represent the overall regime and variability associated with that regime. | Gove et al. 2013, Wedding et al. 2018 |
| Irradiance | Solar radiation at the ocean surface from 4 km MODIS, 8-day composites from 2002-2013. Only long-term mean was retained as other metrics were correlated with each other and metrics of temperature. Long-term mean was most closely associated with our hypothesis of the effects of irradiance on reef conditions. | Gove et al. 2013, Wedding et al. 2018 |
| Waves | Wave power from 0.5-1 km hourly data from the Simulating Waves Nearshore (SWAN) model from 2000-2013. Anomaly frequency and anomaly maximum were retained given they represent different processes and were uncorrelated with each other. | Gove et al. 2013, Wedding et al. 2018 |
| Chl-*a* | Chlorophyll-*a* from 4 km MODIS, 8-day composites from 2002-2013. Anomaly maximum and frequency, and long-term mean were retained as they represent distinct processes and were not correlated. | Gove et al. 2013, Wedding et al. 2018 |
| Habitat | | |
| Depth | Blended depth from in situ surveys, 1999-2001 LiDAR surveys conducted by the Army Corps of Engineers (SHOALS), LiDAR surveys conducted by the Army Corps in 2013 (CZMIL), and aerial imaging spectroscopy data provided by Arizona State University's Global Airborne Observatory (Asner et al. 2020). | (see methods) |
| Rugosity | Blended slope of slope from 1999-2001 LiDAR surveys conducted by the Army Corps of Engineers (SHOALS), LiDAR surveys conducted by the Army Corps in 2013 (CZMIL), and imaging spectroscopy data provided by Arizona State University's Global Airborne Observatory (Asner et al. 2020). | (see methods) |
| Habitat type | Four categories of habitat types: reef (coral dominated hardbottom), pavement, boulder, and other hardbottom based on vector-based maps of habitat produced from aerial imagery by NOAA's Biogeography Branch. | Battista et al. 2007 |

Table S5. Gelman-Rubin statistics for each herbivore driver model and the benthic model assessing convergence of MCMC. Values near one indicate convergence.

| | | Total Herbivore | | Browser | | Grazer | | Scraper | |
|---|---|---|---|---|---|---|---|---|---|
| **Parameter set** | **N** | **Min** | **Max** | **Min** | **Max** | **Min** | **Max** | **Min** | **Max** |
| Beta | 28 | 1.00 | 1.03 | 1.00 | 1.01 | 1.00 | 1.03 | 1.00 | 1.01 |
| Intercept - dataset | 7 | 1.00 | 1.01 | 1.00 | 1.01 | 1.00 | 1.02 | 1.00 | 1.01 |
| Intercept - moku | 40 | 1.00 | 1.03 | 1.00 | 1.02 | 1.00 | 1.03 | 1.00 | 1.01 |
| Intercept - year | 11 | 1.00 | 1.01 | 1.00 | 1.01 | 1.00 | 1.00 | 1.00 | 1.01 |

| | | Calcified:Fleshy | |
|---|---|---|---|
| **Parameter set** | **N** | **Min** | **Max** |
| Beta | 23 | 1.00 | 1.02 |
| Intercept - dataset | 6 | 1.00 | 1.00 |
| Intercept - moku | 37 | 1.00 | 1.02 |
| Intercept - year | 11 | 1.00 | 1.00 |

### Supplemental Methods

#### *Depth and rugosity predictors*

Depth information was missing from *in situ* observations for 20% of the data, and rugosity was not collected by most programs nor was it available in a comparable way by those that did collect metrics of rugosity. Remotely-sensed bathymetric data is reliable for depth and can be used to calculate rugosity. Bathymetric data from 1999-2001 aerial LiDAR surveys conducted by the Army Corps of Engineers (SHOALS) is the most comprehensive dataset available but has significant gaps that precluded statewide estimates. Thus, we combined those data with two other data sources: aerial LiDAR surveys conducted by the Army Corps in 2013 (CZMIL) and aerial imaging spectroscopy-derived depth data at 2 m resolution collected by Arizona State University's Global Airborne Observatory (GAO) [1]. SHOALS and CZMIL data were processed from point cloud data to depth rasters at 5 m and 2 m resolution, respectively. From each of these three depth surfaces, rugosity, measured as topographic rugosity or 'slope of slope', was derived on a 3x3 pixel neighborhood at the native resolution, and the mean value for depth and rugosity were calculated within a 60 m radius of each pixel. To derive a singular depth metric from these three remotely sensed datasets, we used a Bayesian linear model with wide Normal priors (mean of zero, precision of 1/1000) on the slopes and an intercept constrained to zero to predict depth as a weighted mean of the three bathymetry datasets. We modeled *in situ* depth (as recorded by divers during underwater surveys) as a response variable and all three remotely sensed bathymetry datasets as predictors. We allowed for points missing *in situ* depth to be drawn from a Uniform prior between zero and 35, 36, or 22 based on the maximum depths for SHOALS, CZMIL, and GAO respectively to predict depth at every site. For rugosity, we developed a similar Bayesian linear model as described for depth, but we could not rely on diver-measured rugosity as our response variable. Instead, we used the CZMIL 2013 data as the response variable and the SHOALS and GAO data sources as predictors. We allowed for points with missing rugosity for SHOALS and or GAO to be drawn from a Uniform prior bounded between zero and 60 to predict rugosity for all sites.

#### *Predictor maps*

To produce predicted maps, each of the 27 predictor surfaces were processed to a consistent 100 m spatial resolution, grid alignment, extent, and projected coordinate system (NAD 83, UTM zone 4). Predictor layers with native resolutions coarser than 100 m were resampled with the nearest neighbor technique in order to avoid introducing artificial data values and to maintain the spatial patterns of the native resolution. Composite depth and rugosity layers were produced at 100 m resolution by aggregating the high resolution (2 m and 5 m) rasters of 60 m radius mean values to 100 m resolution by calculating the mean of pixels within 100 m x 100 m blocks. For the composite depth surface, raster cell values were filled by priority rank order of data sources, with CZMIL having highest priority, followed by SHOALS, and GAO data was then only used areas where no LiDAR exists. For rugosity, we used a similar priority-based method with CZMIL first but then transformed/weighted values for SHOALS and GAO using a linear predictive relationship between each data source and CZMIL data.

##### *Categorizing soft bottom*

We considered pixels as dominated by soft bottom when the dominant class was 'Sand/Mud' in the vector-based habitat maps produced by NOAA's Biogeography Branch [2], and also further scrutinized a small subset of pixels defined as 'unknown' data by overlaying remotely sensed percent cover of sand estimated at 2 m resolution from [3] and characterized a pixel as soft bottom when the average cover of sand at 100 m resolution was greater than or equal to 85%.

##### *Interpretation of predicted maps*

The uncertainty in our estimates at the 100 m resolution reflects the uncertainty in the effect of each driver on the response variable, and at the moku scale the uncertainty also reflects the spatial variability of the drivers. However, there are other sources of uncertainty and variation that we did not account for, including uncertainty in the predictor values and unexplained variance. The models assume that the driver variables are measured exactly – that is, that there is no error in the observed value of the driver data or when extrapolated to the 100 m map resolution. The models also assume that the driver value at the 100 m resolution applies uniformly at that scale. Further, our models include driver variables and hierarchical spatial variation at the moku scale. These models explain only a fraction of the variability in the observed data. The remaining, unexplained variance is attributable to uncertainty in drivers, additional driver variables that were not included in the model, and fundamental stochasticity in the observation process.

##### *Supplemental methods literature cited*


1. Asner GP, Vaughn NR, Balzotti C, Brodrick PG, Heckler J. 2020 High-Resolution Reef Bathymetry and Coral Habitat Complexity from Airborne Imaging Spectroscopy. *Remote Sens.* **12**, 310.
2. Battista TA, Costa BM, Anderson SM. 2007 Shallow-water benthic habitats of the main eight Hawaiian Islands (DVD). *NOAA Tech. Memo. NOS NCCOS* **61**.
3. Asner GP, Vaughn NR, Heckler J, Knapp DE, Balzotti C, Shafron E, Martin RE, Neilson BJ, Gove JM. 2020 Large-scale mapping of live corals to guide reef conservation. *Proc. Natl. Acad. Sci.* **117**, 33711–33718.
